# Robust inference of positive selection on regulatory sequences in human brain

**DOI:** 10.1101/2020.03.09.984047

**Authors:** Jialin Liu, Marc Robinson-Rechavi

## Abstract

A long standing hypothesis is that divergence between humans and chimpanzees might have been driven more by regulatory level adaptions than by protein sequence adaptations. This has especially been suggested for regulatory adaptions in the evolution of the human brain. There is some support for this hypothesis, but it has been limited by the lack of a reliable and powerful way to detect positive selection on regulatory sequences. We present a new method to detect positive selection on transcription factor binding sites, based on Orr’s sign test applied to a machine learning model of binding. Unlike other methods, this requires neither defining a priori neutral sites, nor detecting accelerated evolution, thus removing major sources of bias. The method is validated in flies, mice, and primates, by a McDonald-Kreitman-like measure of polymorphism vs. divergence, by experimental binding site gains and losses, and by changes in expression levels. We scanned the signals of positive selection for CTCF binding sites in 29 human and 11 mouse tissues or cell types. We found that human brain related cell types have the highest proportion of positive selection. This is consistent with the importance of adaptive evolution on gene regulation in the evolution of the human brain.

## Introduction

It has long been suggested that changes in gene regulation have played an important role in human evolution, and especially in the evolution of the human brain and behaviour (King and Wilson 1975; Anon 2005). Many human and chimpanzee divergent traits (Varki and Altheide 2005) cannot be explained by protein sequence adaptations. For example, there is little evidence to link protein sequence adaptations to traits related to cognitive abilities (Franchini and Pollard 2015). Conversely there is some evidence of brain-specific gene expression divergence in humans (Enard et al. 2002), which is consistent with a role of regulatory evolution. Yet a central question remains open: which regulatory changes were adaptive, if any? A major limitation in answering this is the lack of a robust model of neutral vs. adaptive evolution for regulatory elements.

One approach to detect adaptive evolution on regulatory elements is to detect noncoding regions with lineage-specific accelerated evolutionary rates (Pollard et al. 2006; Prabhakar et al. 2006; Gittelman et al. 2015). For example, Gittelman et al. (2015) found human accelerated regions close to genes annotated to terms such as brain or neuron development. A major caveat is that such acceleration may result from neutral mechanisms such as biased gene conversion (Galtier and Duret 2007) rather than from selection. A second approach is to use a MK test framework (McDonald and Kreitman 1991; Ludwig and Kreitman 1995; Andolfatto 2005; Arbiza et al. 2013; Gronau et al. 2013). This approach has two limitations. First, an expected neutral divergence to polymorphism ratio needs to be defined, whereas defining neutral sites for regulatory elements is difficult and can bias results (Zhen and Andolfatto 2012). And second it lacks power on individual elements, since many regulatory elements are short and present very few variable sites (Andolfatto 2005).

We have developed a new method to detect adaptive evolution of transcription factor binding sites (TFBSs), based on predicted binding affinity changes. As a proof-of-principle, we first applied this method to well-known transcription factors, such as CEBPA and CTCF, in species triples focused on human, mouse, or fly. We validated it with three independent lines of evidence: our evidence of positive selection is associated to higher empirical binding affinity, higher substitution to polymorphism ratio in sequence, and lower variance in expression of neighbouring genes. Then, we used this method to detect positive selection of CTCF binding sites in 29 human tissues or cell types. We found the highest positive selection in brain samples, followed by male reproductive system. The same analysis in mouse found the highest positive selection in the digestive system, with no special signal in the brain. Thus, we provide clear evidence for adaptive evolution of gene regulation in the human brain.

## Results

### Detecting positive selection on transcription factor biding sites

We propose a computational model to detect positive selection on transcription factor binding sites (TFBSs), or any other elements for which we have experimental evidence similar to ChIP-seq (Figure 1, and Methods). Briefly, a gapped k-mer support vector machine (gkm-SVM) classifier is trained on ChIP-seq peaks (here, TFBSs). This allows to compute SVM weights of all possible 10-mers, which are predictions of their contribution to transcription factor binding affinity (Lee et al. 2015). We can then predict the binding affinity impact of substitutions by calculating deltaSVM, the difference of sum weights between two homologous sequences. We compare each empirical TFBS to an ancestral sequence inferred from alignment with a sister species and an outgroup.

**Figure 1:**
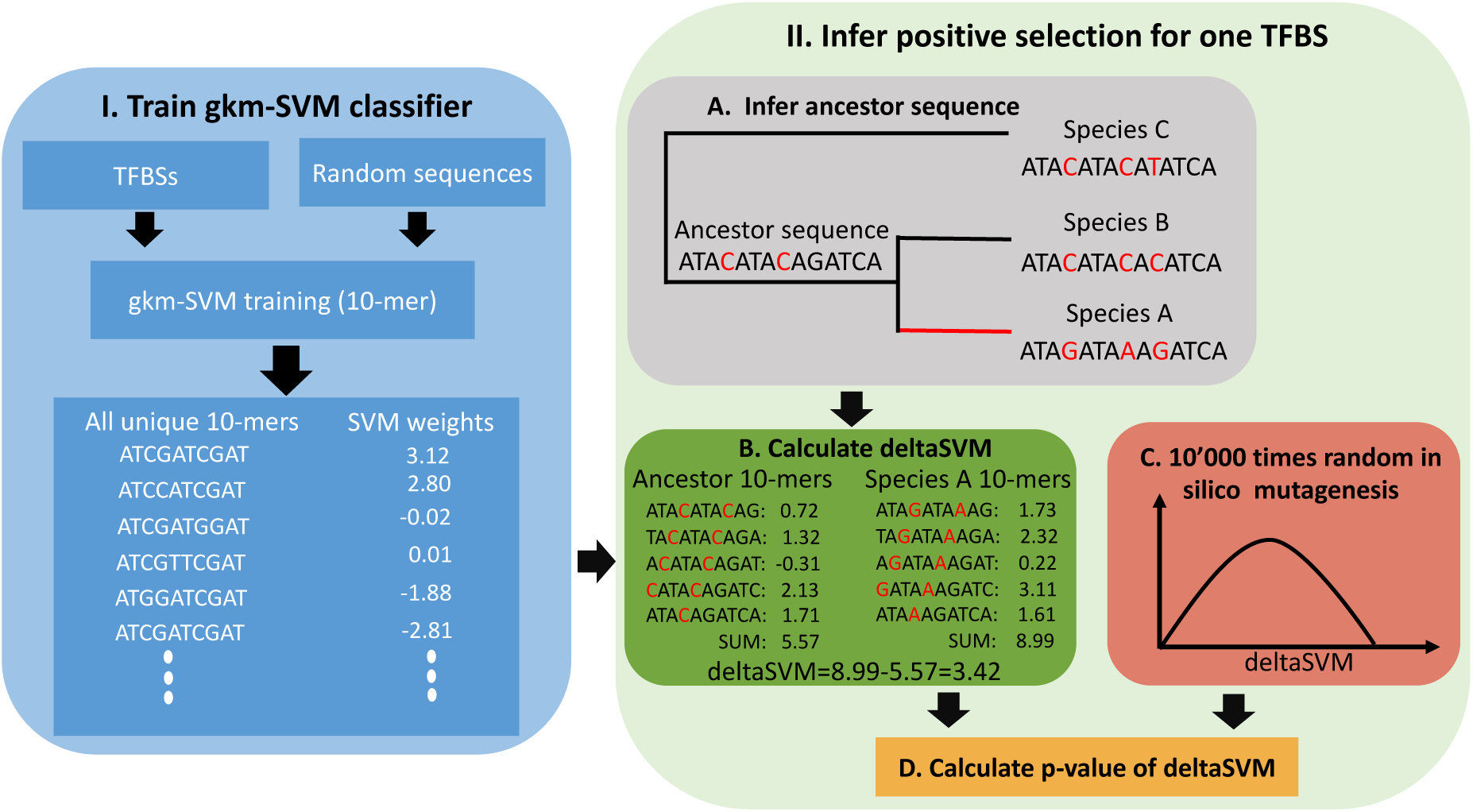
Illustration of the procedure for inferring positive selection. The method includes two parts. Part I (left) is the gapped k-mer support vector machine (gkm-SVM) model training. The gkm-SVM classifier was trained by using TFBSs as a positive training set and randomly sampled sequences from the genome as a negative training set. Then, SVM weights of all possible 10-mers, the contributions of prediction transcription factor binding affinity, were generated from the gkm-SVM. Part II (right) part is the positive selection inference. The ancestor sequence was inferred from sequence alignment with a sister species (species B) and an outgroup (species C). Then, the binding affinity change (deltaSVM) of the two substitutions accumulated in the red branch leading to species A were calculated based on the weight list. The significance of the observed deltaSVM was evaluated by comparing it with a null distribution of deltaSVM, constructed by scoring the same number of random substitutions 10000 times.

Adaptive evolution on TFBSs is expected to push them from a sub-optimal towards an optimal binding strength, or from an old optimum to a new one (e.g. in response to changing environment). Thus TFBSs evolving adaptively are expected to accumulate substitutions which consistently change the phenotype to stronger or to weaker binding, whereas TFBSs evolving under purifying selection are expected to accumulate substitutions which increase or diminish binding in approximately equal measure, around a constant optimum. This reasoning follows the principle of Orr’s sign test of phenotypes (Orr 1998). In practice, this should lead to a large absolute value of deltaSVM under adaptive selection. We estimate by randomisation a p-value specific to each individual TFBS and to its number of substitutions (see Methods). Thus, our method can infer the action of natural selection pushing a TFBS to a new fitness peak of either higher (positive deltaSVM) or lower (negative deltaSVM) binding affinity than its ancestral state.

### Detecting positive selection on liver TFBSs in *Mus musculus*

We first applied our method to a large set of TFBSs in the liver of three mouse species (*Mus musculus domesticus* C57BL/6J, *Mus musculus castaneus* CAST/EiJ, and *Mus spretus* SPRET/EiJ), identified by ChIP-seq for three liver-specific transcription factors, CEBPA, FOXA1, and HNF4A (Stefflova et al. 2013). We inferred positive selection on the lineage leading to C57BL/6J after divergence from CAST/EiJ (Figure 2A). For the sake of simplicity, we only present the results of CEBPA in the main text; results are consistent for FOXA1 and HNF4A (Supplementary materials). We first trained a gkm-SVM on 41945 CEBPA binding sites in C57BL/6J (see Methods). The gkm-SVM very accurately separates CEBPA binding sites and random sequences (Figure 2B). Based on the experimental ChIP-seq peaks in the three species, using SPRET/EiJ as an outgroup, we identified three categories of CEBPA binding sites: conserved in all three species (“conserved”, 24280 sites), lineage specific gain in C57BL/6J (“gain”, 6304 sites), and lineage specific loss in C57BL/6J (“loss”, 6692 sites). Based on whole genome pairwise alignments of C57BL/6J to CAST/EiJ and to SPRET/EiJ, we derived the substitutions accumulated on the C57BL/6J lineage for each CEBPA binding site (see Methods). We only kept binding sites with at least two substitutions, leading to 5114, 1445, and 1497 TFBSs for conserved, gain, and loss categories respectively. For each binding site, we calculated a deltaSVM value, and inferred its significance by random in silico mutagenesis (see Methods).

**Figure 2:**
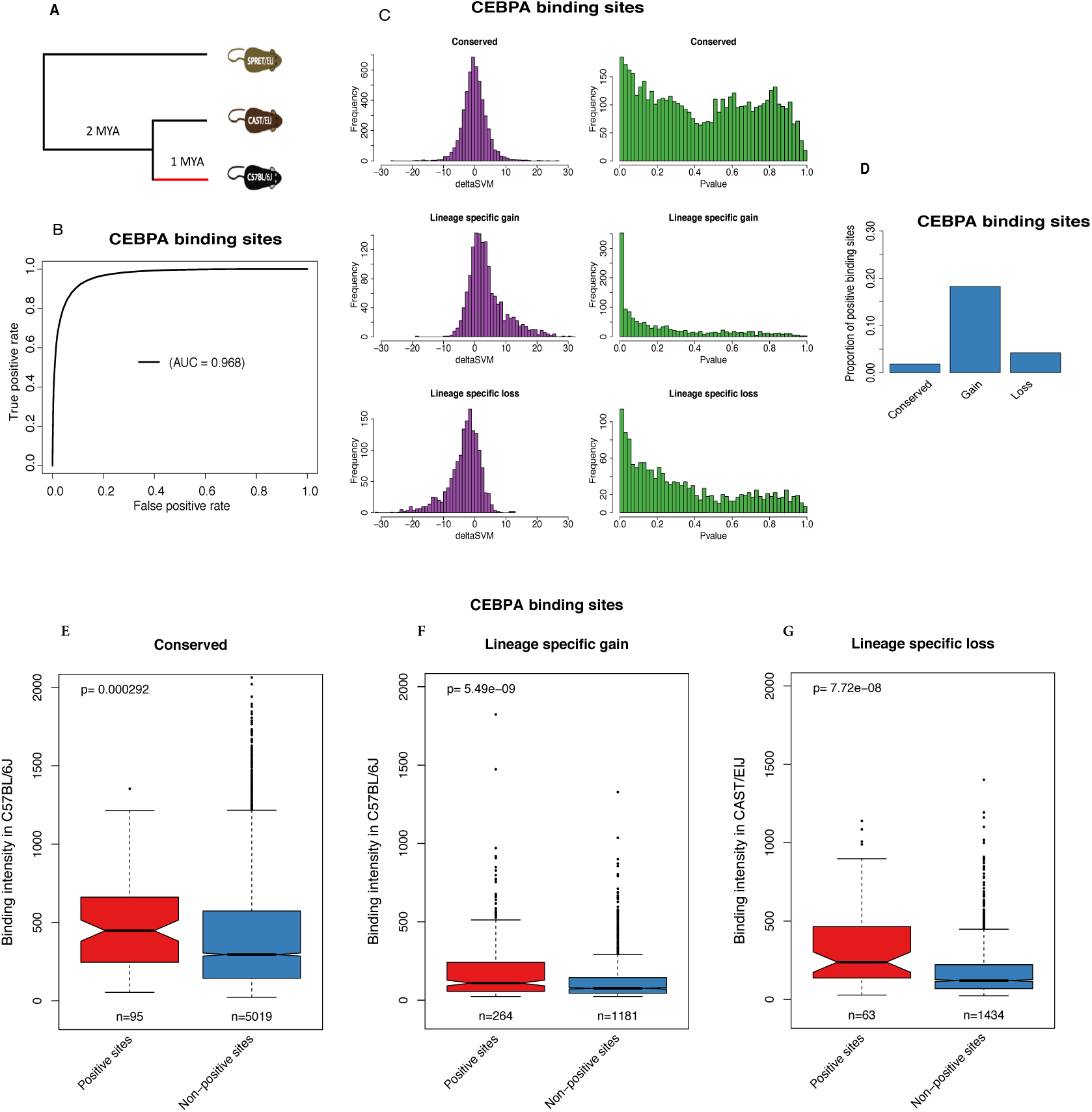
Mouse CEBPA biding sites study. A. Topological illustration of the phylogenetic relationships between the three mouse species used to detect positive selection on CEBPA binding sites. We want to detect positive selection which occurred on the lineage of C57BL/6J, after divergence from CAST/EiJ, as indicated by the red branch. SPRET/EiJ is the outgroup used to infer binding site sequence in the ancestor of C57BL/6J and CAST/EiJ. B. Receiver operating characteristic (ROC) curve for gkm-SVM classification performance on CEBPA binding sites (5-fold cross validation). The AUC value represents the area under the ROC curve and provides an overall measure of predictive power. C. The left hand graphs are the distributions of deltaSVM for conserved, gain, and loss binding sites. The right hand graphs are the distributions of deltaSVM p-values (test for positive selection) for conserved, gain, and loss binding sites. D. The proportion of CEBPA binding sites with evidence of positive selection in conserved, gain, and loss binding sites. E-G. Comparison of biding intensity between positive sites and non-positive sites for mouse CEBPA. The number of binding sites in each category is indicated below each box. The p-values from a Wilcoxon test comparing categories are reported above boxes. The lower and upper intervals indicated by the dashed lines (“whiskers”) represent 1.5 times the interquartile range, or the maximum (respectively minimum) if no points are beyond 1.5 IQR (default behaviour of the R function boxplot). Positive sites are binding sites with evidence of positive selection (deltaSVM p-value < 0.01), non-positive sites are binding sites without evidence of positive selection. E. Conserved binding sites. F. Lineage specific gain binding sites. G. Lineage specific loss binding sites. We compare the binding intensity from CAST/EiJ, as an approximation for ancestral binding intensity, between positive loss binding sites and non-positive loss binding sites.

We plot the distributions of deltaSVMs and of their corresponding p-values for each binding site evolutionary category (Figure 2C). As expected, the distribution of deltaSVMs is symmetric for conserved, has a skew towards positive deltaSVMs for gain, and a skew towards negative deltaSVMs for loss. These results confirm that the gkm-SVM based approach can accurately predict the effect of substitutions on transcription factor binding affinity change. For the distribution of p-values, in all binding site categories, there is a skew of p-values near zero, indicating some signal of positive selection. Gain has the most skewed distribution of p-values towards zero. Hereafter we will use 0.01 as a significant threshold to define positive selection, but results are robust to different thresholds (see **Validation based on ChIP-seq binding intensity**). This identifies almost 20% of gain having evolved under positive selection (Figure 2D), relative to 4% of loss, and 2% of conserved. Random substitutions tend to decrease the binding affinity rather than increase it (Figure S1), because it’s easier to break a function than to improve it. Thus our method could be biased towards reporting as positive sites with more left shifted null distributions. However, this is not the case (Figure S2).

In summary, we found widespread positive selection driving the gain of CEBPA binding sites. We also found some evidence of positive selection driving loss, or increase of binding affinity in some conserved sites. For the other two transcription factors (FOXA1 and HNF4A), we found very consistent patterns (Figure S3, S4).

### Validation based on ChIP-seq binding intensity

We expect that conserved or gained sites which evolved under positive selection with positive deltaSVM should have increased binding affinity. Thus the positive binding sites (PBSs) should have higher biding affinity than non-positive selection binding sites (non-PBSs) in the focal species C57BL/6J. This is indeed the case (Figure 2E and 2F). In addition, conserved TFBSs have higher activity than recently evolved ones (‘gain’). For loss, however, the PBSs have a strong decrease in binding affinity, so we expect higher binding affinity of PBS in the ancestor. Using CAST/EiJ as an approximation for ancestor binding affinity, this is indeed the case (Figure 2G). Results are also consistent with different p-value thresholds (Figure S5). We performed the same validations in FOXA1 and HNF4A, with consistent results (Figure S6).

### Validating the inference of positive selection with human liver TFBSs

To further validate our method, we took advantage of the abundant population genomics transcriptomics data in humans. We inferred positive selection of CEBPA binding sites in the human lineage after divergence from chimpanzee, with gorilla as outgroup (Figure 3A). As in mouse, the gkm-SVM trained from 15806 CEBPA binding sites in human can very accurately separate TFBSs and random sequences (Figure 3B). The distribution of deltaSVMs is slightly asymmetric, with a higher proportion of positive values (Figure 3C). This is because these binding sites contain both conserved and gain, but no loss (since we detect only in the focal species). Based on the distribution of p-values, 7.5% of CEBPA binding sites are predicted to have evolved adaptively in the human lineage.

**Figure 3:**
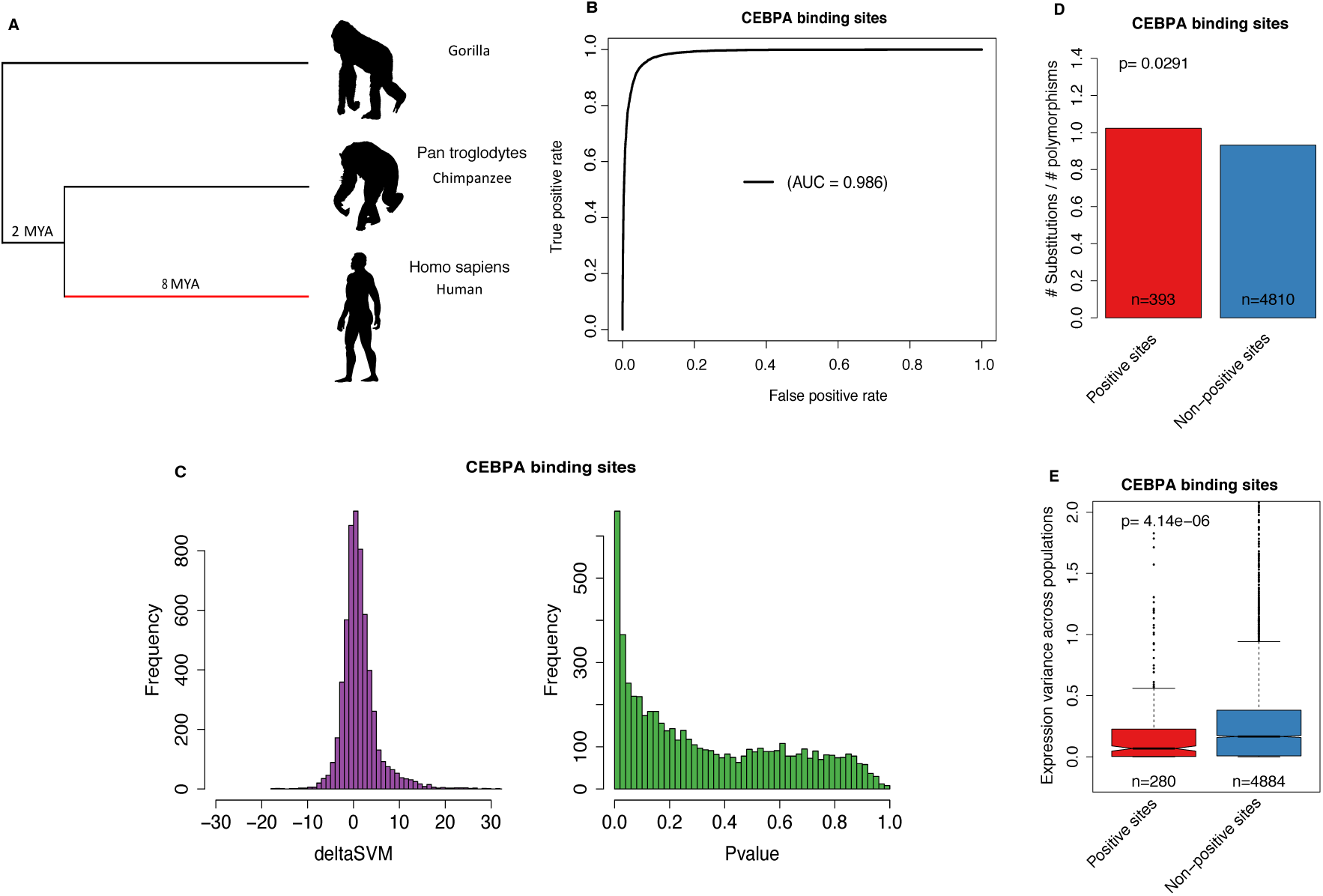
Human CEBPA biding sites study. A. Topological illustration of the phylogenetic relationships between human, chimpanzee and gorilla. We detected positive selection which occurred on the lineage of human after divergence from chimpanzee, as indicated by the red branch. Gorilla is the outgroup used to infer binding site sequence in the ancestor of human and chimpanzee. B. Receiver operating characteristic (ROC) curve for gkm-SVM classification performance on CEBPA binding sites (5-fold cross validation). The AUC value represents the area under the ROC curve and provides an overall measure of predictive power. C. The left graph is the distribution of deltaSVM. The right graph is the distribution of deltaSVM p-values (test for positive selection). D. The ratio between the number of substitutions and the number of polymorphisms (SNPs) for CEBPA binding sites. Positive sites are binding sites with evidence of positive selection (deltaSVM p-value < 0.01), non-positive sites are binding sites without evidence of positive selection. The p-value from Fisher’s exact test is reported above the bars. E. Comparison of expression variance of putative target genes (closest gene to a TFBS) between positive sites and non-positive sites. The number of binding sites in each category is indicated below each box. The p-values from a Wilcoxon test comparing categories are reported above boxes. The lower and upper intervals indicated by the dashed lines (“whiskers”) represent 1.5 times the interquartile range, or the maximum (respectively minimum) if no points are beyond 1.5 IQR (default behaviour of the R function boxplot). Positive sites are binding sites with evidence of positive selection (deltaSVM p-value < 0.01), non-positive sites are binding sites without evidence of positive selection.

Using the MK framework (McDonald and Kreitman 1991), we predict that PBSs should have higher substitutions to polymorphisms ratios than non-PBSs. Note that we do not need to define neutral sites a priori. As expected, we found that the PBSs have a significantly higher ratio of fixed nucleotide changes between human and chimpanzee to polymorphic sites in human than non PBSs (Figure 3D). This is an external validation that our method detects positive selection, as the input did not contain any information about polymorphism.

Besides a higher substitutions to polymorphisms ratio, we also expect that the expression of PBSs putative target genes (see Methods) should be more conserved among human populations. If the expression of PBSs target genes is an adaptive trait in humans, further changes in expression will reduce fitness. Moreover, recent adaptive sweeps are expected to have reduced variability for the regulation of these genes. As expected, we found that PBSs target genes have significantly lower expression variance across human populations than non-PBSs target genes (Figure 3E).

Thus, results from different sources of information support the expectations of our PBS predictions. We performed the same analyses in HNF4A, and results are consistent (Figure S7). These results strongly suggest that our method is detecting real adaptive evolution signals.

### Detecting positive selection of TFBSs in *Drosophila melanogaster*

By using a MK test framework (McDonald and Kreitman 1991), Ni et al. (2012) detected signatures of adaptive evolution on CTCF binding sites in *D. melanogaster*. They reported that positive selection has shaped CTCF binding evolution, and that newly gained binding sites show a stronger signal of positive selection than conserved sites. We applied our method to the same data as used in Ni et al. (2012). We detected positive selection in the *D. melanogaster* lineage after divergence from *D. simulans* (Figure S16A and S16B). Consistent with the findings of Ni et al. (2012), we observed widespread positive selection for both conserved and gain (Figure S16C). In addition, the gain has a higher proportion of positive selection than conserved (Figure S16D). As Ni et al. (2012) did not report specific sites, we cannot compare results more precisely. For lineage specific loss binding sites, however, we did not detect any signal of positive selection (Figure S16C). Interestingly, the proportion of positive selection in *D. melanogaster* is much higher than in *Mus musculus*. For example, we find almost 40% of gain under positive selection in *D. melanogaster*, twice the proportion in *Mus musculus*. It should be noted that different transcription factors and tissues were used, which complicates direct comparison.

### Adaptive evolution of CTCF binding sites across tissues in human

To test whether there is stronger adaptive evolution of gene regulation in some human tissues, we applied our method to 80074 CTCF binding sites across 29 adult tissues or primary cell types (hereafter “cell types”; see Table S2). We chose CTCF because it was the factor with the largest number of tissues or primary cell types studied in a consistent manner, by the ENCODE consortium (The ENCODE Project Consortium 2012; Davis et al. 2018). CTCF is well known as a transcriptional repressor, but it also involved in transcriptional insulation and chromatin architecture remodelling (Phillips and Corces 2009). The gkm-SVM model trained from one cell type can accurately predict the biding sites in another cell type, and the model trained with all CTCF binding sites has better performance than the model trained with cell type specific binding sites (Figure S8). Thus we used a general gkm-SVM rather than different models for different cell types.

We detected 3.52% of positive binding sites (PBSs) for adaptation on the human lineage (Figure S9A). We found that PBSs have higher substitutions to polymorphisms ratio than non-PBSs (Figure S10). In addition, PBSs are associated with a lower number of active cell types (Figure S11A) than non-PBSs, consistent with the prediction that pleiotropy can limit adaptive evolution (Wagner and Zhang 2011). We ranked cell types according to the proportion of binding sites that exhibit statistically significant evidence of positive selection. Brain related cell types have a higher proportion of positive selection than other cell types (Figure 4A). This pattern is consistent if we only use tissue specific CTCF binding sites (Figure S12A). Choroid plexus epithelial cell, brain microvascular endothelial cell, and retinal pigment epithelial cell have notably high PBS frequencies. Non brain related nervous system cell types do not share this high positive selection. Nor does in vitro differentiated neural cell, which may reflect that they do not preserve the signal of specific in vivo differentiated cells. Notably, these brain related cell types also have a higher fraction of substitutions fixed by positive selection (see Methods) than other cell types, except lower leg skin (Figure S13).

**Figure 4:**
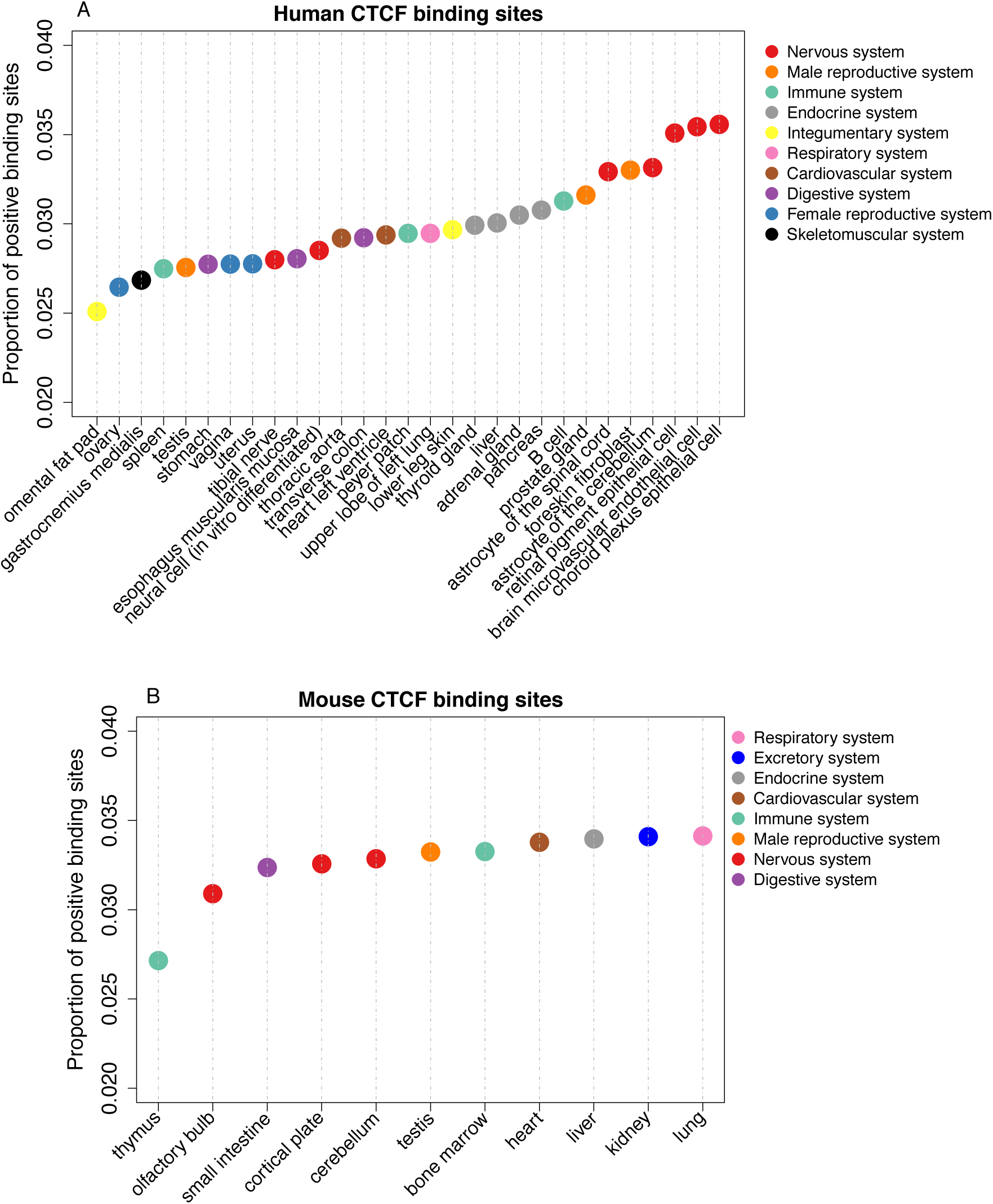
Proportion of positive CTCF binding sites in different tissues or cell types. Positive binding sites are binding sites with evidence of positive selection (deltaSVM p-value < 0.01). Colours correspond to broad anatomical systems. A. CTCF binding sites in 29 human tissues or cell types. B. CTCF binding sites in 11 mouse tissues.

To test whether the high regulatory adaptive evolution in brain is general to mammals, we performed the same analysis on CTCF binding data from 11 mouse adult tissues (Table S2; Figure S14). We investigated adaptive evolution in the *M. musculus* branch after divergence from *M. spretus*, a similar evolutionary divergence as that between human and chimpanzee (Enard et al. 2002). Similarly to human, we detected 3.54% binding sites which evolved under positive selection (Figure S9B) and found PBSs associated with a lower number of active cell types (Figure S11B). However, no tissue type had especially high adaptive evolution, and brain related tissues were among the lowest (Figure 4B). When restricting to tissue specific CTCF binding sites, lung has notably high adaptive evolution (Figure S12B).

## Discussion

### A robust test for positive selection on regulatory elements

Detecting positive selection on regulatory sequences has long been a difficult problem (Zhen and Andolfatto 2012). Nearby non-coding regions are often used as a neutral reference (Andolfatto 2005; Haygood et al. 2007; Arbiza et al. 2013), but such neutrality is difficult to establish. Our approach does not require defining a priori neutral sites, but instead considers the effects of variation on activity (Berg et al. 2004; Moses 2009; Smith et al. 2013). Moreover, positive selection on a background of negative selection might not elevate the evolutionary rate above the neutral expectation, yet consistent changes in binding affinity can still be detectable. Indeed, the TFBSs of cell types detected under selection do not necessarily evolve faster (Figure S15). In principle, our method can also be applied to other genomic regions for which experimental peaks are available, such as open chromatin regions or histone modification regions.

Because positive selection on regulatory sequences is difficult to determine, it is important to validate our predictions with independent evidence. The most important validation is that predictions made independently of population data verify the expectations of higher substitution to polymorphism ratio (Figure 3D). Both this and the lower expression variance of neighbouring genes (Figure 3E) are consistent with the prediction that positive selection will increase divergence but remove polymorphism (McDonald and Kreitman 1991), and that recently selected phenotypes will be under stronger purifying selection. Moreover, binding affinity change occurs in the direction predicted by our model (Figure 2E-G), and we can verify the prediction that pleiotropy limits adaptation (Wagner and Zhang 2011) (Figure S11).

Despite its advantages, our method can still be improved. For example, in the null model of sequence evolution, we assume independent mutation patterns at each base-pair site and a uniform mutation rate over all sites. But both mutation rate and pattern can depend on neighbouring nucleotides (Krawczak et al. 1998). These limitations of our null model might explain why the observed p-values do not quite follow the expected uniform distribution for high values.

### The importance of regulatory adaptation on human brain evolution

Our results support the long proposed importance of adaptive regulatory changes in human brain evolution (King and Wilson 1975). They are remarkably consistent with accelerated gene expression evolution in the human brain, but neither in human blood or liver, nor in rodents, from Enard et al. (2002). Previous studies on human regulatory sequence evolution reported acceleration in brain related functions, but could not demonstrate adaptive evolution nor direct activity in the brain (Enard et al. 2002; Pollard et al. 2006; Prabhakar et al. 2006; Haygood et al. 2007; Gittelman et al. 2015). The reported link between human accelerated regions and function was very indirect, depending both on the attribution of a region to the closest gene, and on the functional annotation of that gene.

The brain related cell types for which we detect a high proportion of positive selection are functionally related with cognitive abilities. For example, for astrocyte, abnormal astrocytic signalling can cause synaptic and network imbalances, leading to cognitive impairment (Santello et al. 2019). In addition, for choroid plexus epithelial cell, its atrophy has been reported to be related with Alzheimer disease (Kaur et al. 2016).

While we did not find a similar pattern by applying the same analysis to mouse, it isn’t possible yet to conclude to a human or primate specific pattern. Indeed, the mouse analyses have two potential caveats. First, for the olfactory bulb and cortical plate in the mouse analyses, there are no corresponding anatomical structures in the human analyses. It is an open question whether the human olfactory bulb and cortical plate also have high adaptation. Second, the human analyses were based on ChIP-seq at cell type level but the mouse analyses were based on ChIP-seq at tissue level. In mouse, the astrocyte in cerebellum may also have high adaptation like the astrocyte in human, but the signal might be diluted by other cell types in cerebellum.

### Regulatory adaptation differs between tissues

Outside of brain cell types, we found that male reproduction system (prostate and foreskin) has higher adaptive regulatory evolution than female reproduction system (ovary, uterus and vagina). This is consistent with the observation of high adaptive sequence evolution in human male reproduction (Wyckoff et al. 2000; Nielsen et al. 2005), and probably caused by sexual selection related selective pressures, such as sperm competition. However testis has a relatively low proportion of adaptive evolution, similar to ovary. This suggests that the high expression divergence in testis (Brawand et al. 2011) is mainly caused by relaxed purifying selection, maybe due to the role of transcription in testis for ‘transcriptional scanning’ (Xia et al. 2020). Outside of the brain, the top adaptive regulatory evolution systems seem to be the same as found for adaptive protein evolution, i.e. male reproduction, immune and endocrine systems (Clark et al. 2003; Bustamante et al. 2005; Nielsen et al. 2005; Daub et al. 2017). The high fraction of substitutions fixed by positive selection in skin is interesting (Figure S13), since skin is both involved in defence against pathogens, and has evolved specifically in the human branch with loss of fur (Brettmann and de Guzman Strong 2018). The lack of adaptive protein sequence evolution despite high adaptive regulatory evolution might be related to selective pressure on proteins in the brain (Drummond and Wilke 2008; Roux et al. 2017).

## Materials and Methods

### Code and data availability

Data files and analysis scripts are available on GitHub: https://github.com/ljljolinq1010/A-robust-method-for-detecting-positive-selection-on-regulatory-sequences

All data analysed during this study are available from public databases listed under each dataset in the relevant Materials and Methods section.

### Mutagenesis for positive selection

#### 1. Training of the gapped k-mer support vector machine (gkm-SVM)

gkm-SVM is a method for regulatory DNA sequence prediction by using *k*-mer frequencies (Ghandi et al. 2014). For the gkm-SVM training, we followed the approach of Lee et al. (2015). Firstly, we defined a positive training set and its corresponding negative training set. The positive training set is ChIP-seq narrow peaks of transcription factors. The negative training set is an equal number of sequences which randomly sampled from the genome with matched the length, GC content and repeat fraction of the positive training set. This negative training set was generated by using “genNullSeqs”, a function of gkm-SVM R package (Ghandi et al. 2016). Then, we trained a gkm-SVM with default parameters except -l=10 (meaning we use 10-mer as feature to distinguish positive and negative training sets). The classification performance of the trained gkm-SVM was measured by using receiver operating characteristic (ROC) curves with fivefold cross-validation. The gkm-SVM training and cross-validation were achieved by using the “gkmtrain” function of “LS-GKM : a new gkm-SVM software for large-scale datasets” (Lee 2016). For details, please check https://github.com/Dongwon-Lee/lsgkm.

#### 2. Generate SVM weights of all possible 10-mers

The SVM weights of all possible 10-mers were generated by using the “gkmpredict” function of “LS-GKM”. The positive value means increasing binding affinity, the negative value means decreasing binding affinity, the value close to 0 means functionally neutral.

#### 3. Infer ancestor sequence

The ancestor sequence was inferred from sequence alignment with a sister species and an outgroup.

#### 4. Calculate deltaSVM

We calculated the sum of weights of all 10-mers for ancestor sequence and focal sequence respectively. The deltaSVM is the sum weights of focal sequence minus the sum weights of ancestor sequence. The positive deltaSVM indicating substitutions increased the binding affinity in the focal sequence, vice versus.

#### 5. Generate Empirical Null Distribution of deltaSVM

Firstly, we counted the number of substitutions between the ancestor sequence and the focal sequence. Then, we generated a random pseudo-focal sequence by randomly introducing the same number of substitutions to the ancestor sequence. Finally, we calculated the deltaSVM between the pseudo-focal sequence and the ancestor sequence. We repeated the above processes 10000 times to get 10000 expected deltaSVMs.

#### 6. Calculate p-value of deltaSVM

For lineage specific gain TFBSs, the p-value was calculated as the probability that the expected deltaSVM is higher than the observed deltaSVM (higher-tail test). For lineage specific lose TFBSs, the p-value was calculated as the probability that the expected deltaSVM is lower than the observed deltaSVM (lower-tail test). For conserved TFBSs, we primarily focused on selection to increase binding affinity, and thus we performed higher-tail test. The motivation for this is that when we have ChIP-seq data in only one species, which is the most common case, the observed peaks are a mix of conserved and gained sites, and thus very little signal of decrease of binding is expected. The p-value can be interpreted as the probability that the observed deltaSVM could arise by chance.

### Mouse validation analysis

#### 1. ChIP-seq data

The narrow ChIP-seq peaks and their corresponding intensity (normalized read count) datasets of three liver specific transcription factors (CEBPA, FOXA1 and HNF4A) in three mouse species (C57BL/6J, CAST/EiJ, SPRET/EiJ) were downloaded from https://www.ebi.ac.uk/research/flicek/publications/FOG09 (Stefflova et al. 2013, accessed in May, 2018). Peaks were called with SWEMBL (http://www.ebi.ac.uk/~swilder/SWEMBL). To account for both technical and biological variabilities of peak calling, Stefflova et al. (2013) carried out the following approaches. For each transcription factor in each species, they first called three sets of peaks: one for each replicate (replicate peek), and one for a pooled dataset of both replicates (pooled peak). Then, the peaks detected from the pooled dataset were used as a reference to search for overlaps in the two other replicates. When a pooled peak overlapped with both replicate peeks (at least one base pair overlap), it was kept for downstream analyses. For the number of peaks and average peak length, please check Table S1.

#### 2. Peak coordinates transfer

Based on pairwise genome alignments between C57BL/6J and CAST/EiJ or SPRET/EiJ, Stefflova et al. (2013) transferred the coordinates of ChIP-seq peaks in both CAST/EiJ and SPRET/EiJ to its corresponding coordinates in C57BL/6J.

#### 3. Sequence alignment files

The sequence alignment files between C57BL/6J and CAST/EiJ or SPRET/EiJ were downloaded from https://www.ebi.ac.uk/research/flicek/publications/FOG09 (Stefflova et al. 2013, accessed in May, 2018).

#### 4. Define different types of binding sites

##### 1) Conserved binding sites

The conserved binding sites were defined as peaks in C57BL/6J which have overlapping peaks (at least one base pair overlap) in the other two species by genome alignment.

##### 2) Lineage specific gain binding sites

The lineage specific gain binding sites defined as peaks in C57BL/6J with no overlapping peaks (at least one base pair overlap) in the other two species.

##### 3) Lineage specific loss binding sites

The lineage specific loss binding sites defined as peaks in CAST/EiJ which overlapping peaks in SPRET/EiJ but not in C57BL/6J.

### Human validation analysis

#### 1. ChIP-seq data

The narrow ChIP-seq peaks datasets of two liver specific transcription factors (CEBPA and HNF4A) in human were downloaded from https://www.ebi.ac.uk/research/flicek/publications/FOG01 (Schmidt et al. 2010, accessed in October, 2018). Peaks were called with SWEMBL (http://www.ebi.ac.uk/~swilder/SWEMBL). Negligible variation was observed between the individuals in terms of peak calling, so Schmidt et al. (2010) pooled replicates into one dataset for peak calling.

#### 2. Sequence alignment files

The pairwise whole genome alignments between human and chimpanzee or gorilla were downloaded from http://hgdownload.soe.ucsc.edu/downloads.html (accessed in December, 2018).

#### 3. Single nucleotide polymorphism (SNP) data

Over 36 million SNPs for 1,092 individuals sampled from 14 populations worldwide were downloaded from phase I of the 1000 Genomes Project ftp://ftp.1000genomes.ebi.ac.uk/vol1/ftp/phase1/analysis_results/integrated_call_sets/ (1000 Genomes Project Consortium 2012, accesed in December, 2018). As suggested by Luisi et al. (2015), we only used SNPs of a subset of 270 individuals from YRI, CEU, and CHB populations.

#### 4. Liver expression data

The library site normalized expression data of 175 livers was downloaded from downloaded from The Genotype Tissue Expression (GTEx) project https://gtexportal.org/home/ (GTEx Consortium 2017, Release V7, accessed in December, 2018). We further transformed it with log_2_.

#### 5. Putative target genes of TFBSs

We assigned the nearest gene to each TFBS as its putative target gene.

### Fly validation analysis

#### 1. ChIP-seq data

The narrow ChIP-seq peaks of transcription factor CTCF in three drosophila species (*D. melanogaster, D. simulans* and *D. yakuba*) were downloaded from https://www.ncbi.nlm.nih.gov/geo/query/acc.cgi?acc=GSE24449 (Ni et al. 2012, accessed in January, 2019). Peaks were called with QuEST(Valouev et al. 2008) at a False Discovery Rate (FDR) <1%. We obtained 2182, 2197 and 2993 peaks with average length of 243bp, 240bp and 201bp for *D. melanogaster, D. simulans* and *D. yakuba* respectively.

#### 2. Peak coordinates transfer

The peaks identified in *D. simulans* and *D. yakuba* were translated onto *D. melanogaster* coordinates by using pslMap (Zhu et al. 2007).

#### 3. Sequence alignment files

The pairwise whole genome alignments between *D. melanogaster* and *D. simulans* or *D. yakuba* were downloaded from http://hgdownload.soe.ucsc.edu/downloads.html (accessed in January, 2019).

4. Define different types of binding sites. These were defined as in human, using *D. melanogaster* vs. *D. simulans* and *D. yakuba*.

### Human CTCF analysis

#### 1. ChIP-seq data

The narrow ChIP-seq peaks of transcription factor CTCF across 29 tissues or cell types (Table S2) were downloaded from ENCODE (The ENCODE Project Consortium 2012). We did not use ChIP-seq datasets from cell lines, and only kept ChIP-seq datasets from tissues and primary cells. Briefly, peaks were called with MACS (Zhang et al. 2008) separately for each replicate. Irreproducible Discovery Rate (IDR) analysis was then performed (Li et al. 2011). Final peaks are the set of peak calls that pass IDR at a threshold of 2%. Peaks identified in different tissues or cell types were integrated by intersecting all peaks across datasets, with at least one base pair overlap used as the merge criteria. Overall we obtained 118970 merged peaks.

#### 2. Sequence alignment files

The pairwise whole genome alignments between human and chimpanzee or gorilla were downloaded from http://hgdownload.soe.ucsc.edu/downloads.html (accessed in December, 2018).

#### 3. Proportion of substitutions fixed by positive selection

We calculated the Proportion of substitutions fixed by positive selection, a measure of effect size of selection, under the MK test framework (McDonald and Kreitman 1991; Smith and Eyre-Walker 2002):

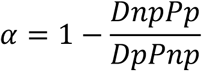

*Dnp* is the substitution number in non-PBSs; *Pp* is the polymorphism number in PBSs; *Dp* is the substitution number in PBSs; *Pnp* is the polymorphism number in non-PBSs.

### Mouse CTCF analysis

#### 1. CHiP-seq data

The narrow ChIP-seq peaks of transcription factor CTCF across 11 tissues (Table S2) were downloaded from ENCODE (The ENCODE Project Consortium 2012). Briefly, peaks were called with MACS (Zhang et al. 2008) separately for each replicate. Irreproducible Discovery Rate (IDR)) analysis was then performed. Final peaks are the set of peak calls that pass IDR at a threshold of 2%. Peak identified in different tissues/cell types were integrated by intersecting all peaks across data sets, with at least 1 base pair overlap used as the merge criteria. Overall we obtained 112657merged peaks.

#### 2. Sequence alignment files

The sequence alignment file between C57BL/6J and SPRET/EiJ, please check “Mouse validation analysis” part of Materials and Methods. The sequence alignment file between C57BL/6J and Caroli/EiJ were downloaded from https://www.ebi.ac.uk/research/flicek/publications/FOG09 (Stefflova et al. 2013, accessed in May, 2018).

## Acknowledgements

We thank Jérôme Goudet, Gunter Wagner, David Garfield, Laurent Duret, and members of the Robinson-Rechavi lab for helpful discussions. Part of the computations were performed at the Vital-IT (http://www.vital-it.ch) Center for high-performance computing of the SIB Swiss Institute of Bioinformatics. JL and MRR are supported by Swiss National Science Foundation grants 31003A_153341 / 1 and 31003A_173048.

## Conflict of Interests

The authors declare that they have no conflict of interest.

## Author contributions

JL designed the work with input from MRR. JL performed data collection and computational analyses. JL and MRR interpreted the results. JL wrote the first draft of the paper. JL and MRR finalized the paper.

## Supplementary figure legends

**Figure S1: The null distribution of deltaSVM for a conserved biding site of mouse CEBPA**.

**Figure S2: Comparison of mean deltaSVM of null distribution between positive sites and non-positive sites for mouse CEBPA**.

For each binding site, we first calculated its mean deltaSVM of null distribution. Then, we compared the mean values between positive sites and non-positive sites. The number of binding sites in each category is indicated below each box. The *p*-values from a Wilcoxon test comparing categories are reported above boxes. The lower and upper intervals indicated by the dashed lines (“whiskers”) represent 1.5 times the interquartile range, or the maximum (respectively minimum) if no points are beyond 1.5 IQR (default behaviour of the R function boxplot). Positive sites are binding sites with evidence of positive selection (deltaSVM *p-*value < 0.01), non-positive sites are binding sites without evidence of positive selection.

**Figure S3: mouse FOXA1 biding sites study** The figure legend is the same as Figure 2A-D.

**Figure S4: mouse HNF4A biding sites study** The figure legend is the same as Figure 2A-D.

**Figure S5: Comparison of biding intensity between positive sites and non-positive sites for mouse CEBPA**.

A-C: Positive sites defined as deltaSVM with *p-*value < 0.05 instead of 0.01 in Figure 2E-G.

D-F: Positive sites defined as deltaSVM with *p-*value < 0.001 instead of 0.01 in Figure 2E-G.

The figure legend is the same as for Figure 2E-G.

**Figure S6: Comparison of biding intensity between positive sites and non-positive sites for mouse FOXA1 and HNF4A**

The figure legend is the same as Figure 2E-G.

**Figure S7: human HNF4A biding sites study** The figure legend is the same as Figure 3.

**Figure S8: Receiver operating characteristic (ROC) curves for gkm-SVM classification performance on human CTCF binding sites**.

AUC values represent areas under the ROC curve and provide an overall measure of predictive power.

A. The results of a 5-fold cross validation on neural CTCF binding sites and matched random sequences.

B. The gkm-SVM trained in B cell used to predict neural CTCF binding sites and matched random sequences.

C. The gkm-SVM trained in all 29 tissues/cell types used to predict neural CTCF binding sites and matched random sequences.

D. The results of a 5-fold cross validation on B cell CTCF binding sites and matched random sequences.

E. The gkm-SVM trained in neural cell used to predict B cell CTCF binding sites and matched random sequences.

F. The gkm-SVM trained in all 29 tissues/cell types used to predict B cell CTCF binding sites and matched random sequences.

**Figure S9: The distribution of deltaSVM and deltaSVM *p*-values for CTCF biding sites in human and mouse**

**Figure S10: Ratio between the number of substitutions and the number of polymorphisms (SNPs) for human CTCF binding sites**.

Positive sites are binding sites with evidence of positive selection (deltaSVM *p-*value < 0.01), non-positive sites are binding sites without evidence of positive selection. The *p*-value from Fisher’s exact test is reported above the bars.

**Figure S11: Comparison of the number of active tissues/cell types for CTCF biding sites between positive sites and non-positive sites**.

The *p*-values from a Wilcoxon test comparing categories are reported above boxes. The lower and upper intervals indicated by the dashed lines (“whiskers”) represent 1.5 times the interquartile range, or the maximum (respectively minimum) if no points are beyond 1.5 IQR (default behaviour of the R function boxplot). Positive sites are binding sites with evidence of positive selection (deltaSVM *p-*value < 0.01), non-positive sites are binding sites without evidence of positive selection.

**Figure S12: Proportion of positive CTCF binding sites in different tissues or cell types**. Here, we only consider cell type or tissue specific CTCF binding sites. Positive binding sites are binding sites with evidence of positive selection (deltaSVM *p-*value < 0.01). Colours correspond to broad anatomical systems.

A. CTCF binding sites in 29 human tissues or cell types.

B. CTCF binding sites in 11 mouse tissues.

**Figure S13: Proportion of substitutions fixed by positive selection in different tissues or cell types**.

**Figure S14: Receiver operating characteristic (ROC) curve for gkm-SVM classification performance on mouse CTCF binding sites**.

The result comes from a 5-fold cross validation on all CTCF binding sites from 11 tissues and matched random sequences. The AUC value represents areas under the ROC curve and provides an overall measure of predictive power.

**Figure S15: Average number of substitutions for non-positive CTCF binding sites in different tissues/cell types**.

Non-positive binding sites are binding sites with evidence of positive selection (deltaSVM *p-*value >= 0.01). The number of substitutions measured by the pairwise whole genome alignments between human and chimpanzee. For each binding site, the number of substitutions was normalized by the length of the binding site.

A. CTCF binding sites in 29 human tissues/cell types.

B. CTCF binding sites in 11 mouse tissues.

**Figure S16: *D. melanogaster* CTCF biding sites study**.

A. Topological illustration of the phylogenetic relationships between the three *Drosophila* species used to detect positive selection on CTCF binding sites. We want to detect positive selection which occurred on the lineage of *D. melanogaster* after divergence from *D. simulans*, as indicated by the red branch. *D. yakuba* is the outgroup used to infer binding site sequence in the ancestor of *D. melanogaster* and *D. simulans*.

B. Receiver operating characteristic (ROC) curve for gkm-SVM classification performance on CEBPA binding sites (5-fold cross validation). The AUC value represents the area under the ROC curve and provides an overall measure of predictive power.

C. The left hand graphs are the distributions of deltaSVM for conserved, gain, and loss binding sites. The right hand graphs are the distributions of deltaSVM *p*-values (test for positive selection) for conserved, gain, and loss binding sites.

D. The proportion of CEBPA binding sites with evidence of positive selection in conserved and gain binding sites.

**Table S1: The number of peaks and average peak length for CEBPA, FOXA1 and HNF4A in three mouse species C57BL/6J, CAST/EiJ and SPRET/EiJ**.

**Table S2: The CTCF ChIP-seq datasets information for human and mouse**.

## Notes

https://github.com/ljljolinq1010/A-robust-method-for-detecting-positive-selection-onregulatory-sequences

### Competing Interest Statement

The authors have declared no competing interest.

### Summary of Updates

Reverted to version 1 upon request of the journal considering the manuscript.

