## Supplementary materials for "Robust inference of positive selection on regulatory sequences in human brain"

### 1 Supplementary figures

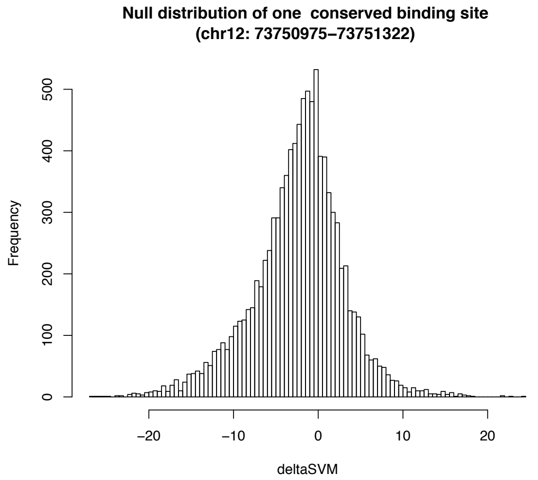


**Figure S1: The null distribution of deltaSVM for a conserved biding site of mouse CEBPA.**

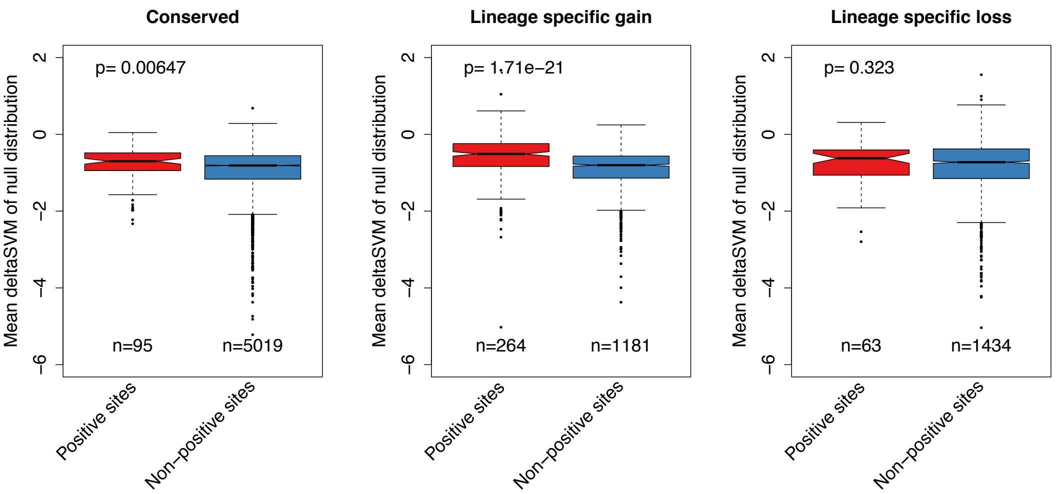


**Figure S2: Comparison of mean deltaSVM of null distribution between positive sites and non-positive sites for mouse CEBPA.**

For each binding site, we first calculated its mean deltaSVM of null distribution. Then, we compared the mean values between positive sites and non-positive sites. The number of binding sites in each category is indicated below each box. The *p*-values from a Wilcoxon test comparing categories are reported above boxes. The lower and upper intervals indicated by the dashed lines ("whiskers") represent 1.5 times the interquartile range, or the maximum (respectively minimum) if no points are beyond 1.5 IQR (default behavior of the R function boxplot). Positive sites are binding sites with evidence of positive selection (deltaSVM *p*-value < 0.01), non-positive sites are binding sites without evidence of positive selection.

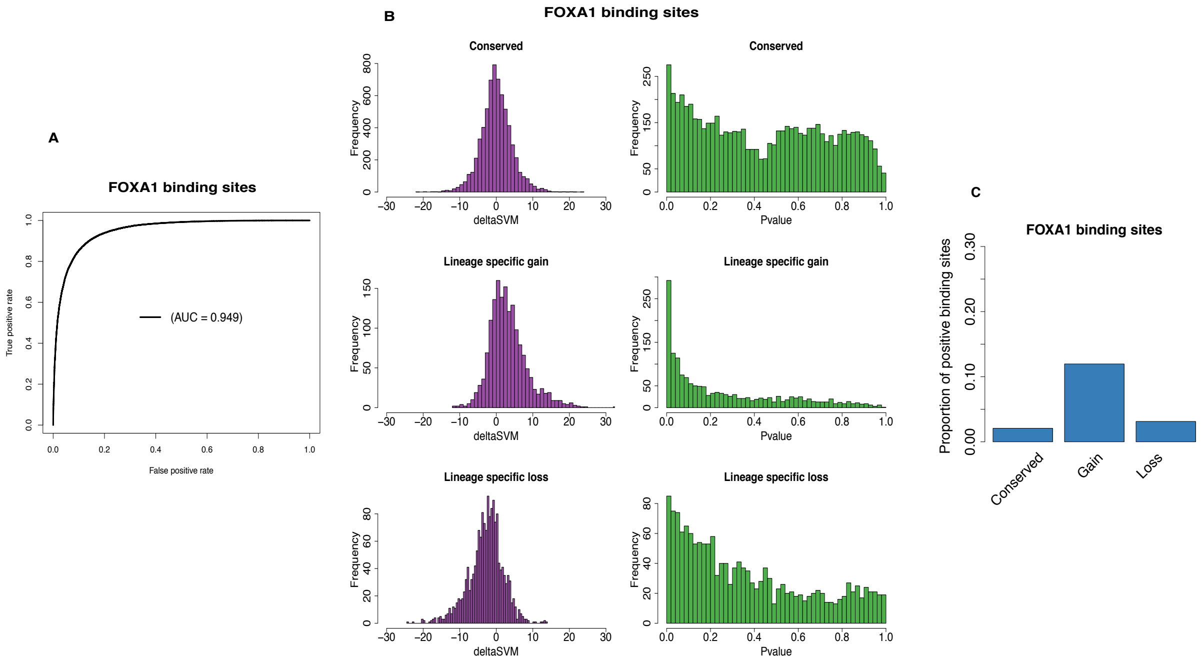


**Figure S3: mouse FOXA1 biding sites study**

The figure legend is the same as Figure 2A-D.

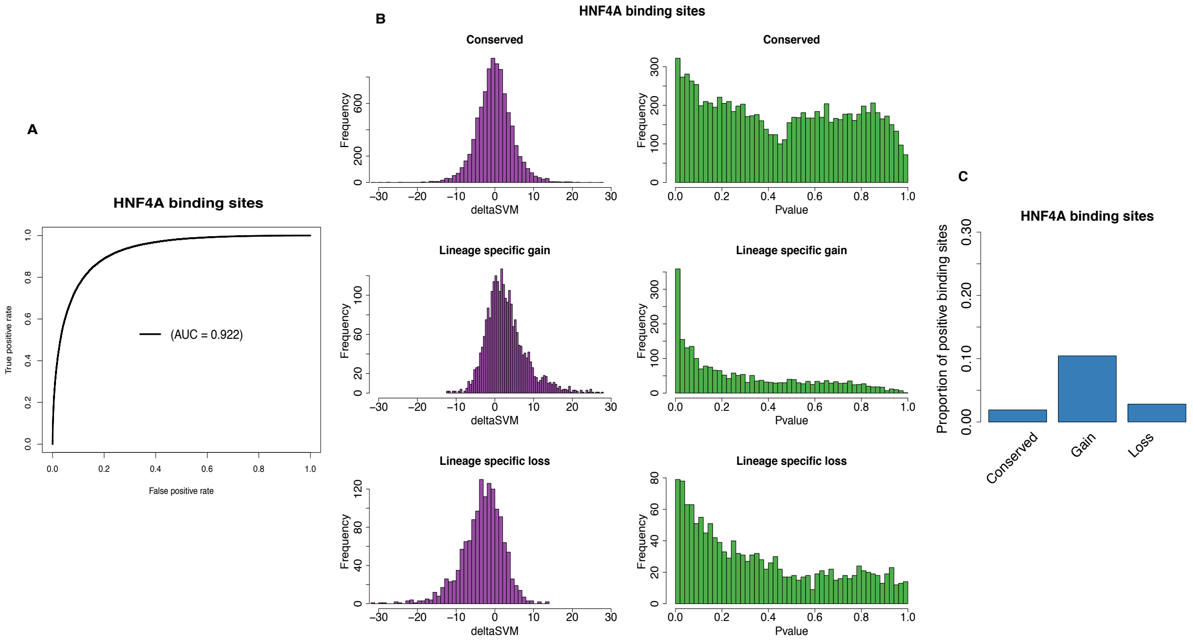


**Figure S4: mouse HNF4A biding sites study**

The figure legend is the same as Figure 2A-D.

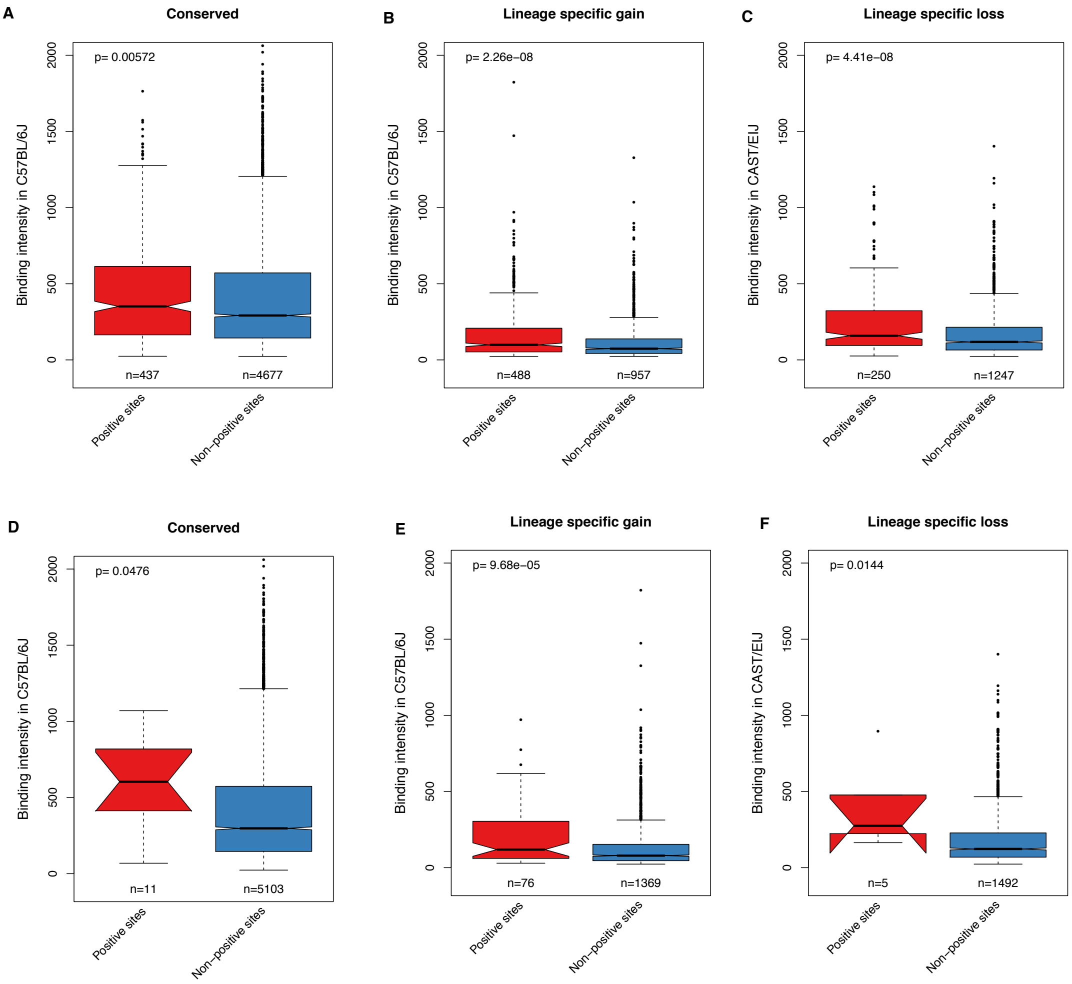


**Figure S5: Comparison of biding intensity between positive sites and non-positive sites for mouse CEBPA.**

A-C: Positive sites defined as deltaSVM with $p$-value < 0.05 instead of 0.01 in Figure 2E-G.

D-F: Positive sites defined as deltaSVM with $p$-value < 0.001 instead of 0.01 in Figure 2E-G.

The figure legend is the same as for Figure 2E-G.

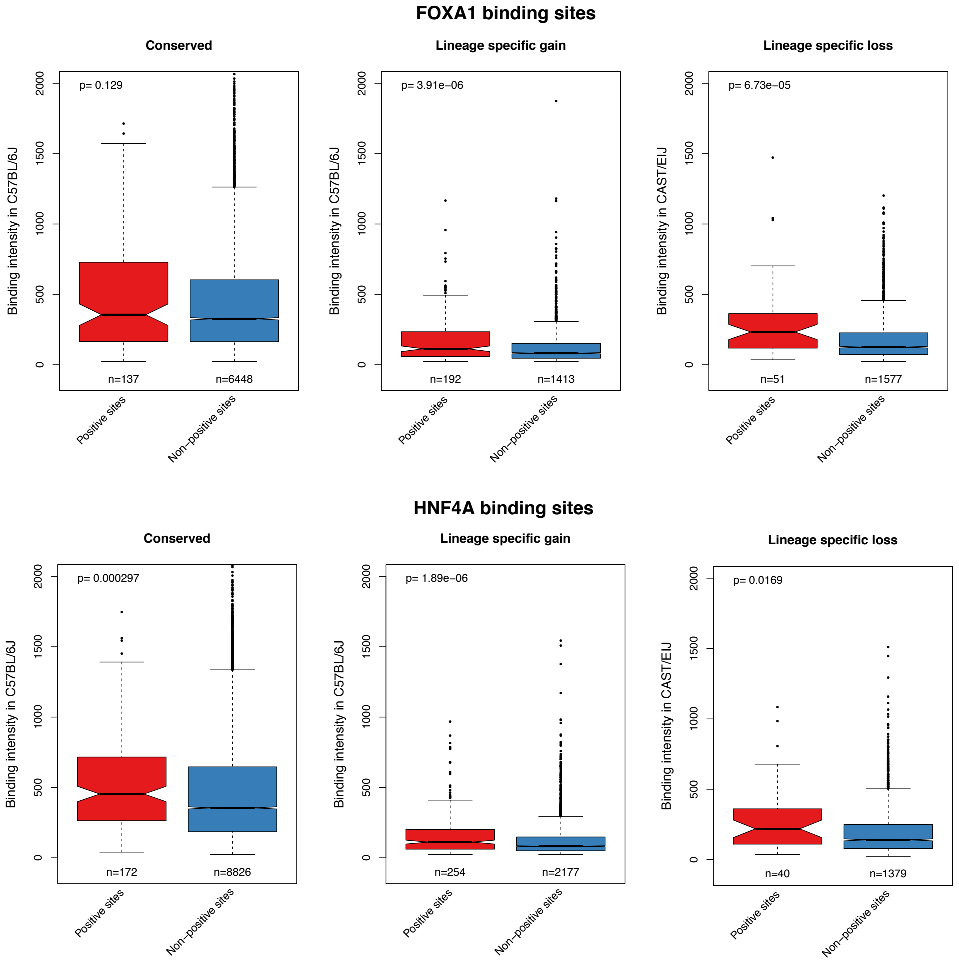


**Figure S6: Comparison of biding intensity between positive sites and non-positive sites for mouse FOXA1 and HNF4A**

The figure legend is the same as Figure 2E-G.

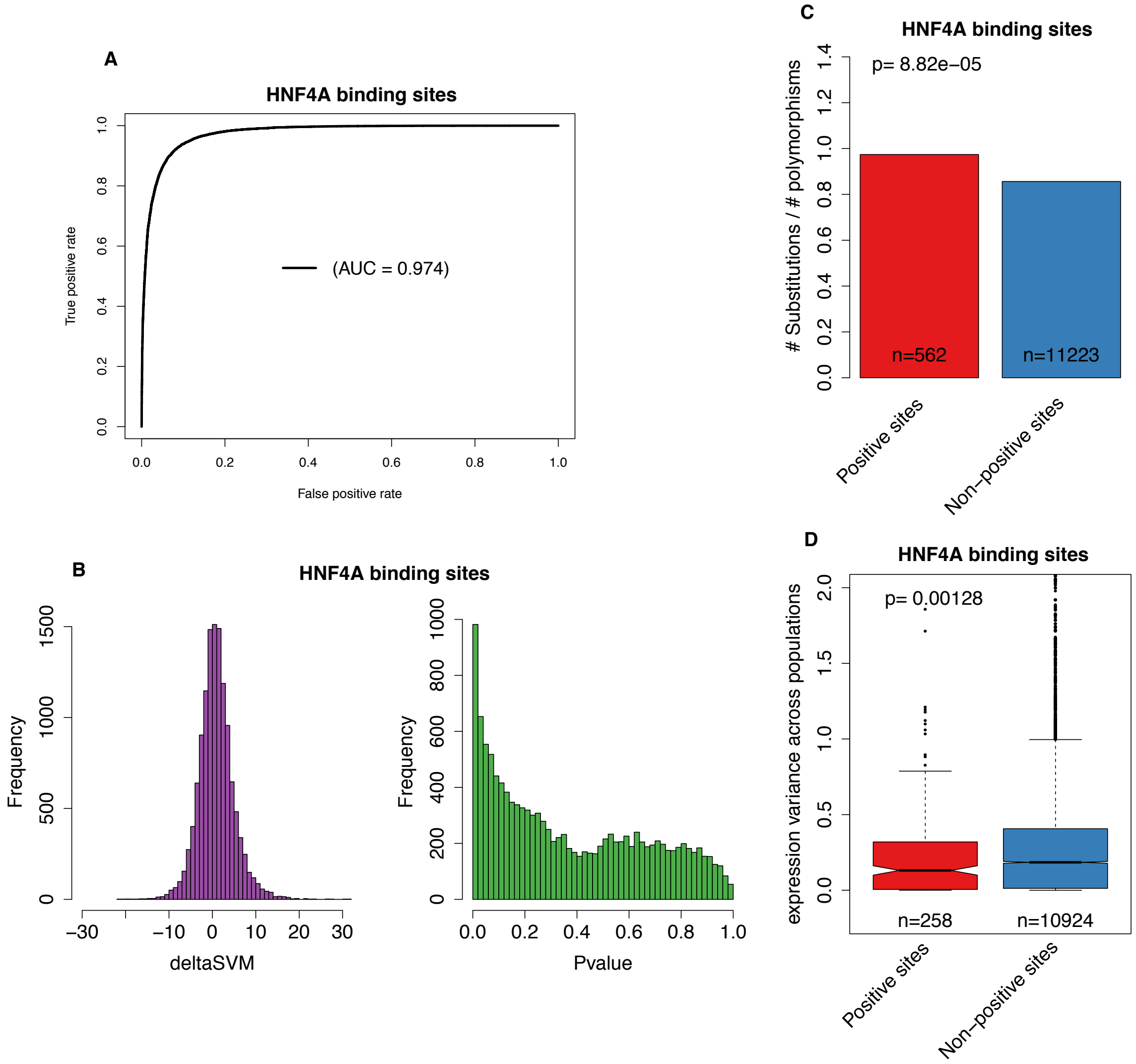


**Figure S7: human HNF4A biding sites study**

The figure legend is the same as Figure 3.

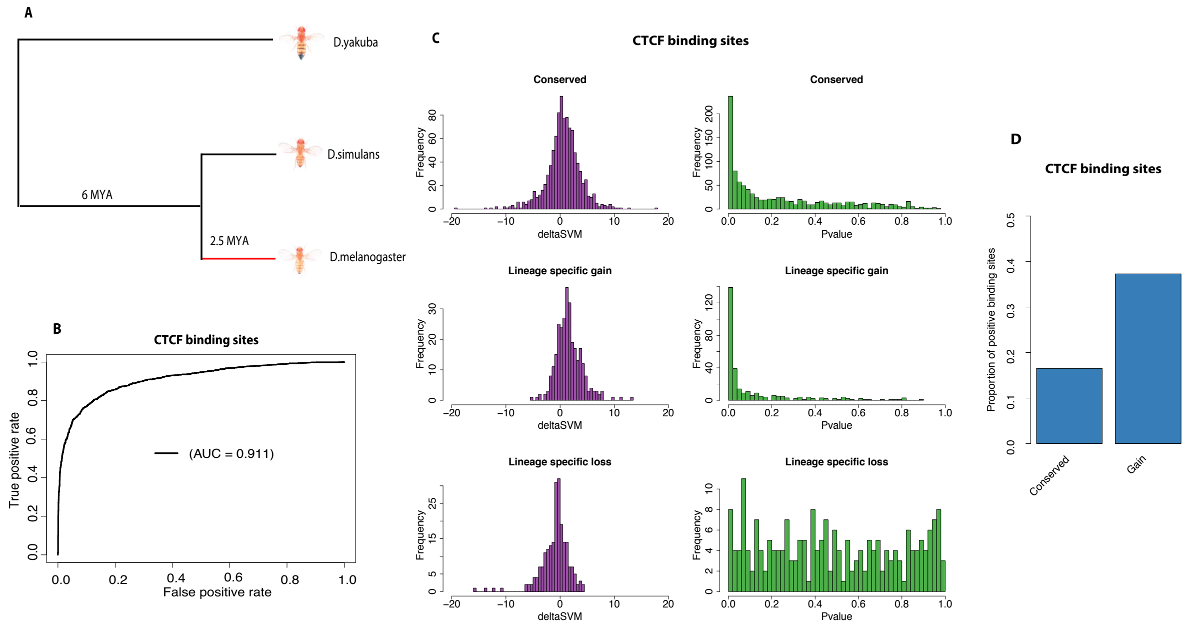


**Figure S8:** ***D. melanogaster*** **CTCF biding sites study.**

A. Topological illustration of the phylogenetic relationships between the three *Drosophila* species used to detect positive selection on CTCF binding sites. We want to detect positive selection which occurred on the lineage of *D. melanogaster* after divergence from *D. simulans*, as indicated by the red branch. *D. yakuba* is the outgroup used to infer binding site sequence in the ancestor of *D. melanogaster* and *D. simulans*.

B. Receiver operating characteristic (ROC) curve for gkm-SVM classification performance on CEBPA binding sites (5-fold cross validation). The AUC value represents the area under the ROC curve and provides an overall measure of predictive power.

C. The left hand graphs are the distributions of deltaSVM for conserved, gain, and loss binding sites. The right hand graphs are the distributions of deltaSVM *p*-values (test for positive selection) for conserved, gain, and loss binding sites.

D. The proportion of CEBPA binding sites with evidence of positive selection in conserved and gain binding sites.

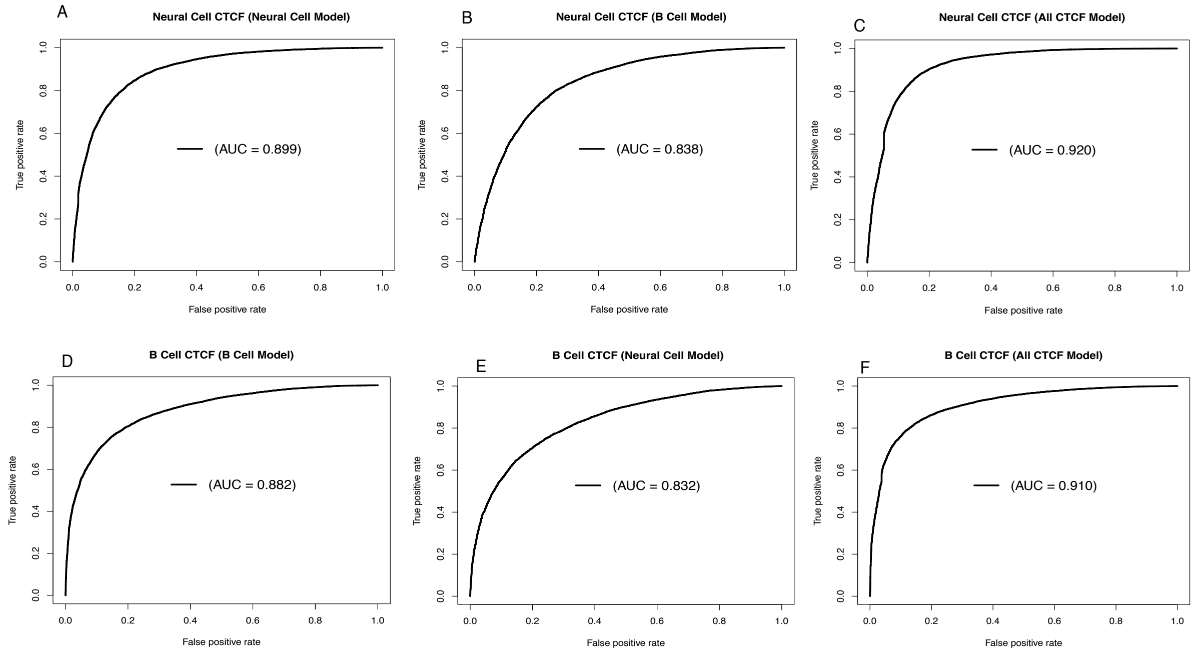


**Figure S9: Receiver operating characteristic (ROC) curves for gkm-SVM classification performance on human CTCF binding sites.**

AUC values represent areas under the ROC curve and provide an overall measure of predictive power.

A. The results of a 5-fold cross validation on neural CTCF binding sites and matched random sequences.

B. The gkm-SVM trained in B cell used to predict neural CTCF binding sites and matched random sequences.

C. The gkm-SVM trained in all 29 tissues/cell types used to predict neural CTCF binding sites and matched random sequences.

D. The results of a 5-fold cross validation on B cell CTCF binding sites and matched random sequences.

E. The gkm-SVM trained in neural cell used to predict B cell CTCF binding sites and matched random sequences.

F. The gkm-SVM trained in all 29 tissues/cell types used to predict B cell CTCF binding sites and matched random sequences.

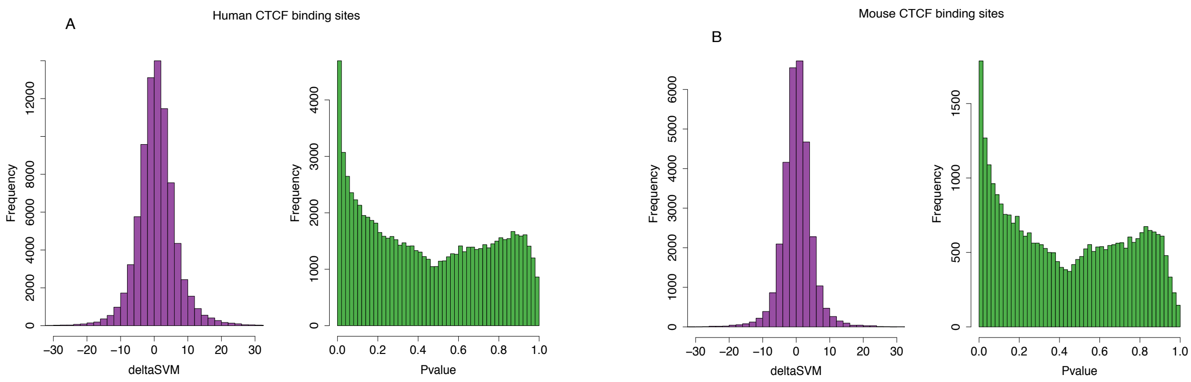


**Figure S10: The distribution of deltaSVM and deltaSVM *p*-values for CTCF biding sites in human and mouse**

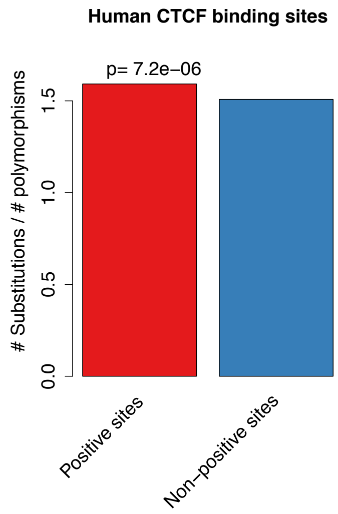


**Figure S11: Ratio between the number of substitutions and the number of polymorphisms (SNPs) for human CTCF binding sites.**

Positive sites are binding sites with evidence of positive selection (deltaSVM *p*-value < 0.01), non-positive sites are binding sites without evidence of positive selection. The *p*-value from Fisher's exact test is reported above the bars.

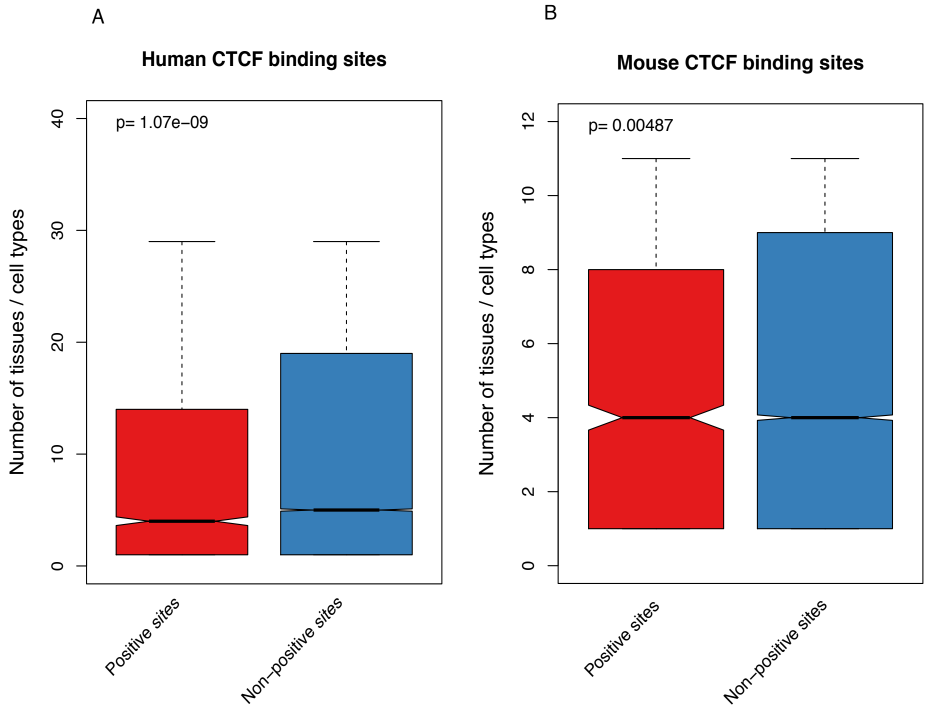


**Figure S12: Comparison of the number of active tissues/cell types for CTCF biding sites between positive sites and non-positive sites.**

The *p*-values from a Wilcoxon test comparing categories are reported above boxes. The lower and upper intervals indicated by the dashed lines ("whiskers") represent 1.5 times the interquartile range, or the maximum (respectively minimum) if no points are beyond 1.5 IQR (default behavior of the R function boxplot). Positive sites are binding sites with evidence of positive selection (deltaSVM *p*-value < 0.01), non-positive sites are binding sites without evidence of positive selection.

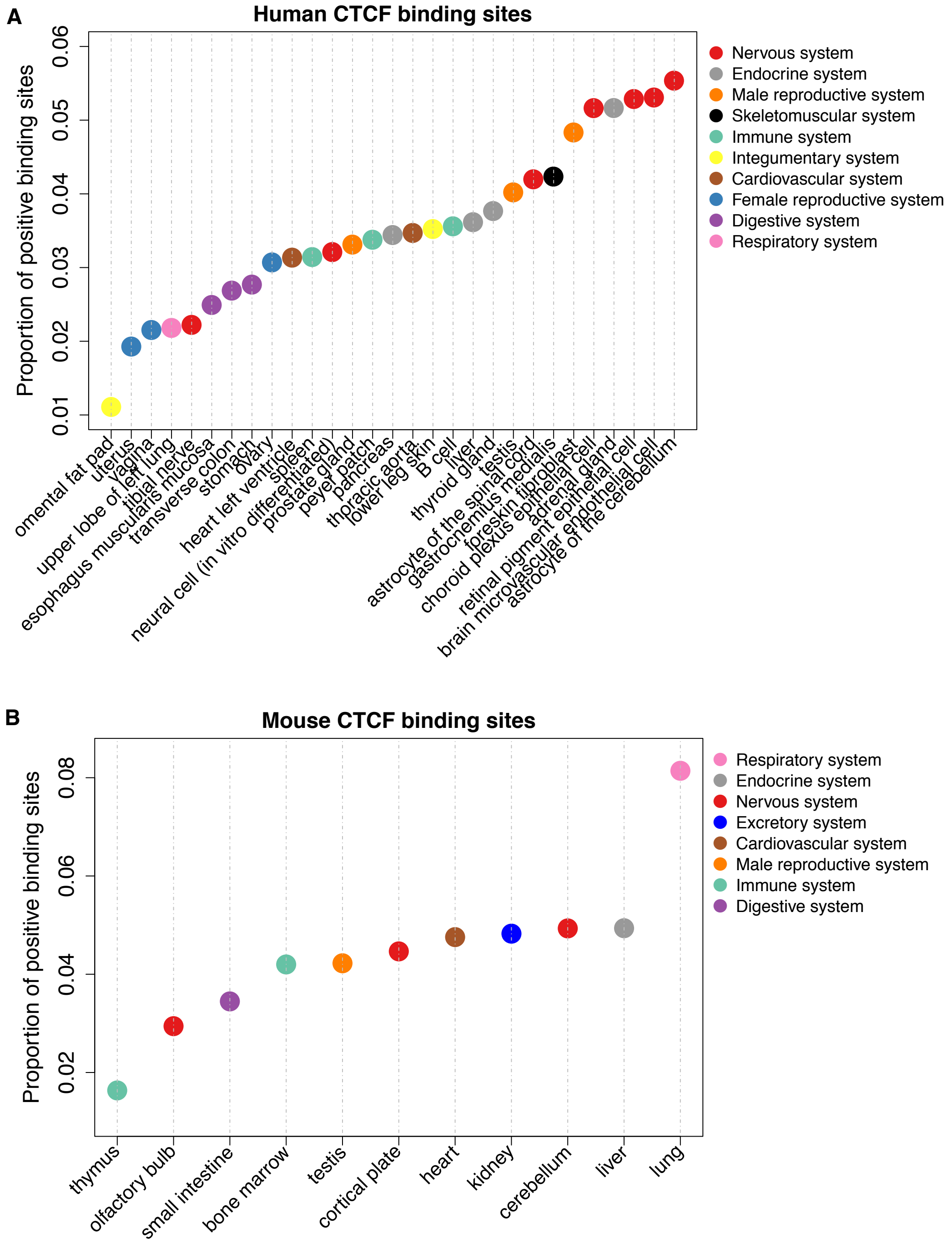


**Figure S13: Proportion of positive CTCF binding sites in different tissues or cell types.** Here, we only consider cell type or tissue specific CTCF binding sites. Positive binding sites are binding sites with evidence of positive selection (deltaSVM $p$-value $< 0.01$). Colors correspond to broad anatomical systems.

A. CTCF binding sites in 29 human tissues or cell types.

B. CTCF binding sites in 11 mouse tissues.

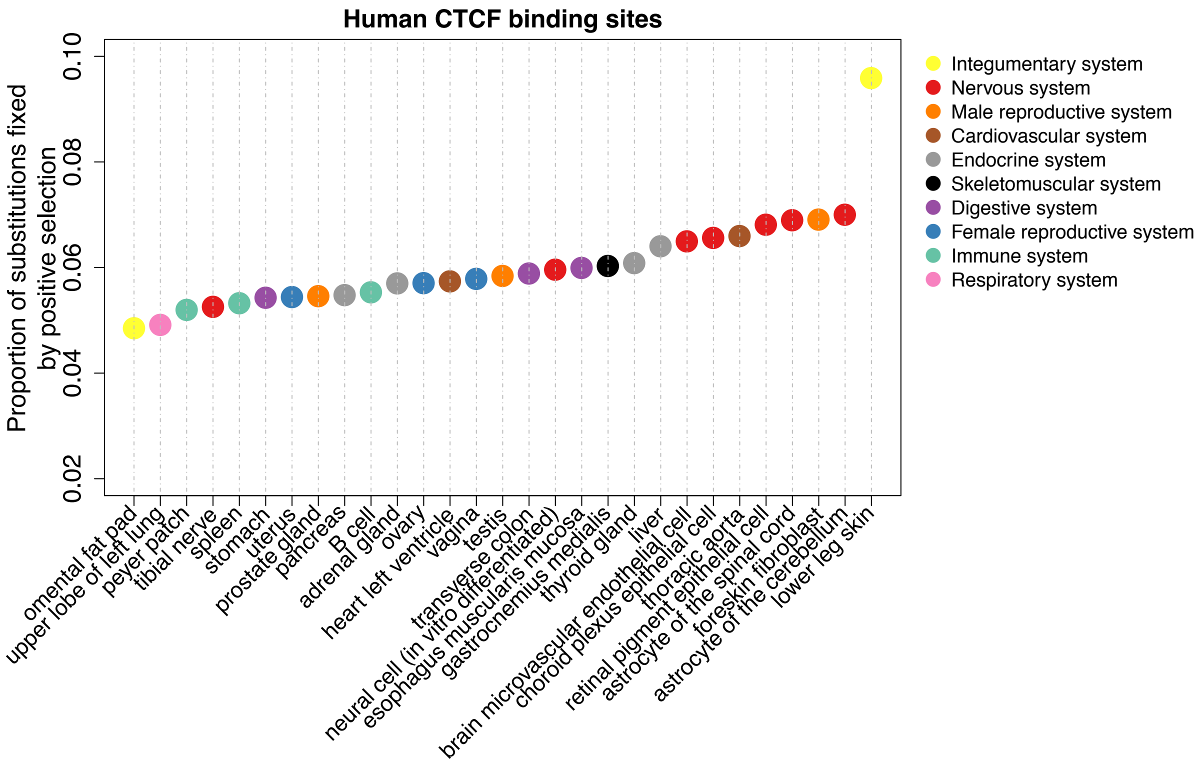


**Figure S14: Proportion of substitutions fixed by positive selection in different tissues or cell types.**

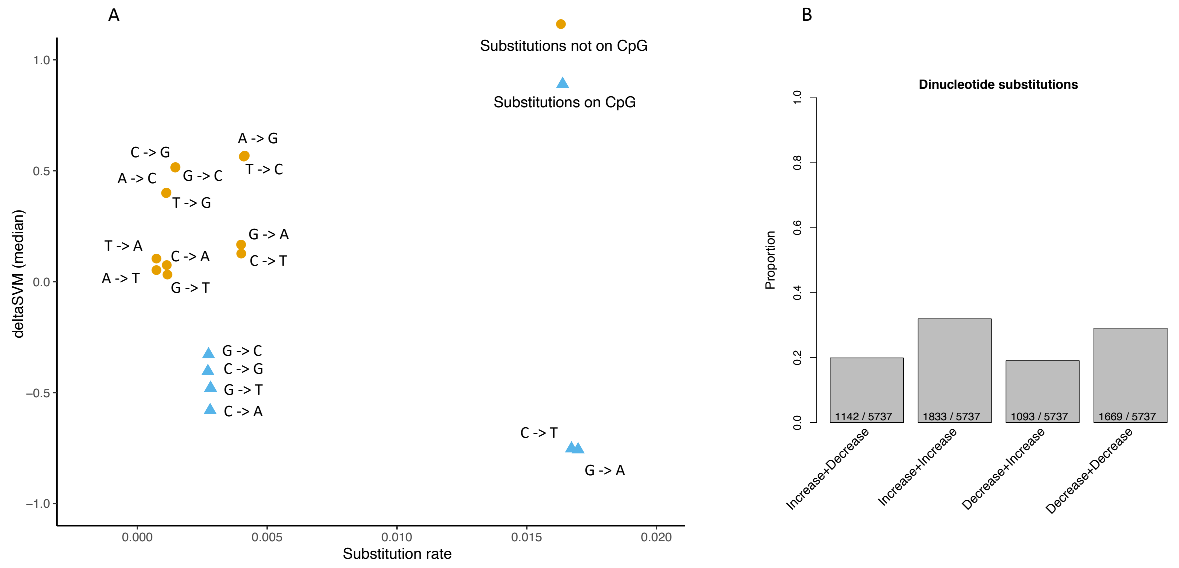


**Figure S15: Substitution type, substitution rate and deltaSVM relationship.**

A. The x-axis is the substitution rate of different types of substitutions, the y-axis is the median deltaSVM of different types of substitutions.

B. The label below the bottom of each bar indicates the direction of binding affinity change of two neighboring substitutions. For example, Increase+Decrease indicates the first substitution increases the binding affinity, but the second substitution decreases the binding affinity. the number of dinucleotide substitutions in each category and the total number of dinucleotide substitutions are indicated in the bottom of each bar.

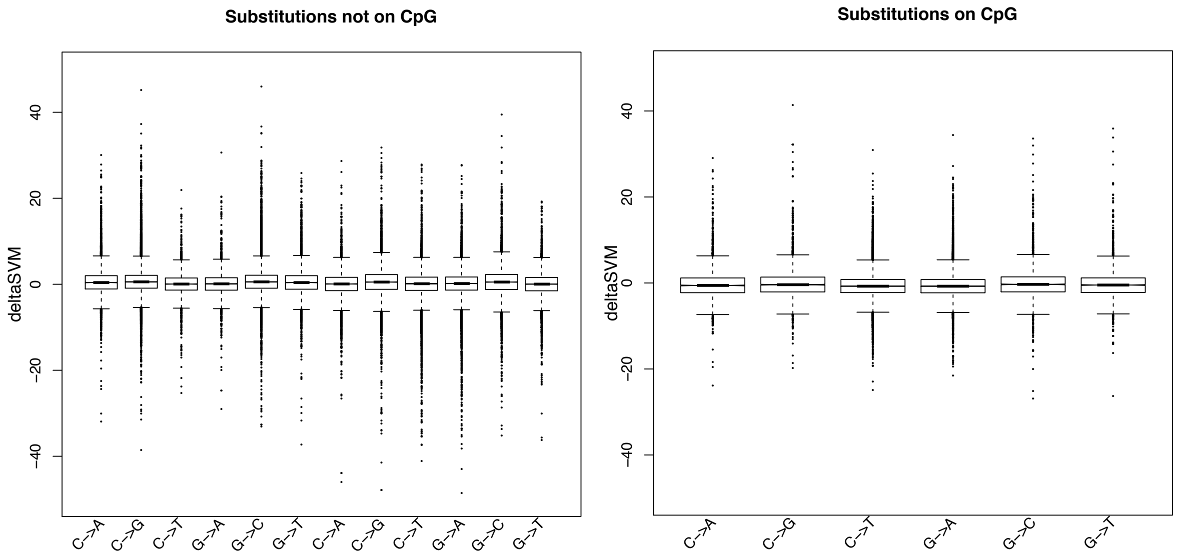


**Figure S16: deltaSVM comparisons between different type of Substitutions.**

The lower and upper intervals indicated by the dashed lines (“whiskers”) represent 1.5 times the interquartile range, or the maximum (respectively minimum) if no points are beyond 1.5 IQR (default behavior of the R function boxplot).

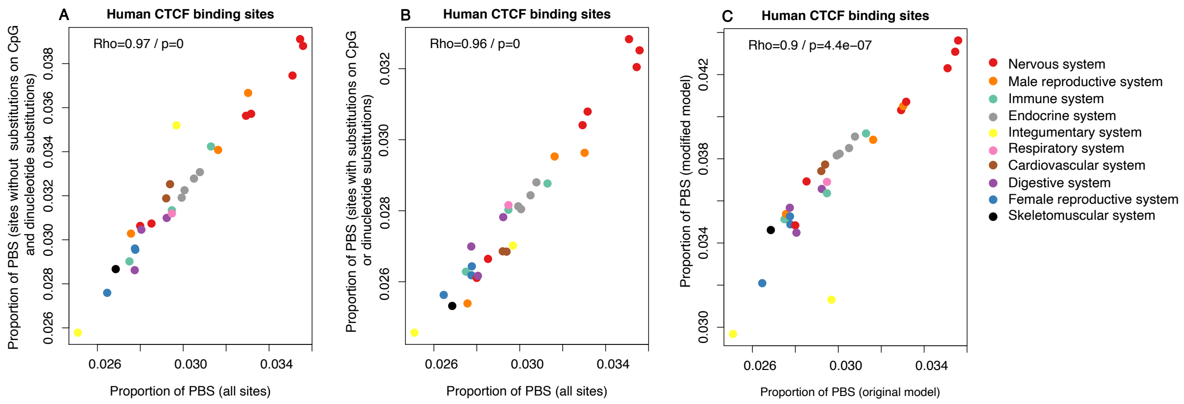


**Figure S17: Correlation of PBS proportion between different analyses.**

PBS, the positive selection sites defined as (deltaSVM *p*-value <0.01).

A. The x axis is the proportion of PBS in different tissues and cell types for all sites, the y axis is the proportion of PBS in different tissues and cell types for sites without substitutions on CpG and without dinucleotide substitutions.

B. x axis is the proportion of PBS in different tissues and cell types for all sites, y axis is the proportion of PBS in different tissues and cell types for sites with substitutions on CpG or with dinucleotide substitutions.

C. x axis is the proportion of PBS in different tissues and cell types from the original analysis, y axis is the proportion of PBS in different tissues and cell types for sites from the new analysis. In the new analysis, we excluded all CpG sequences and dinucleotide substitution sequences, and controlled the transition and transversion rate.

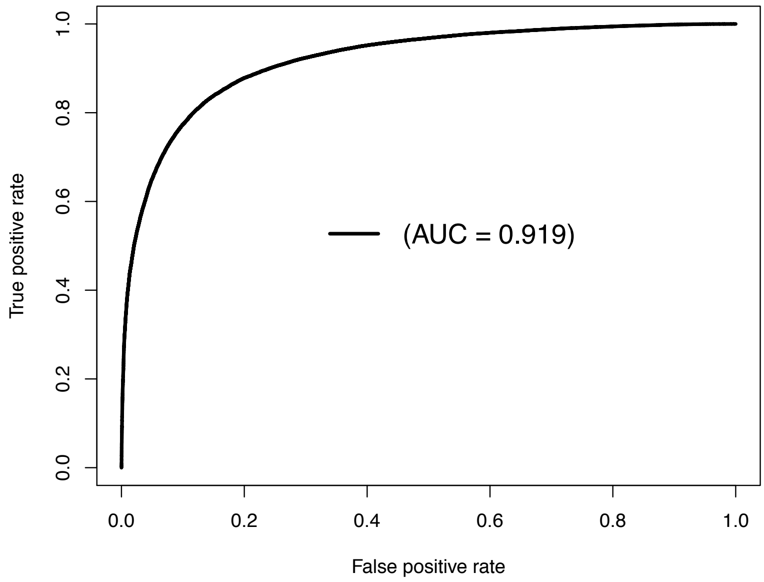


**Figure S18: Receiver operating characteristic (ROC) curve for gkm-SVM classification performance on mouse CTCF binding sites.**

The result comes from a 5-fold cross validation on all CTCF binding sites from 11 tissues and matched random sequences. The AUC value represents areas under the ROC curve and provides an overall measure of predictive power.

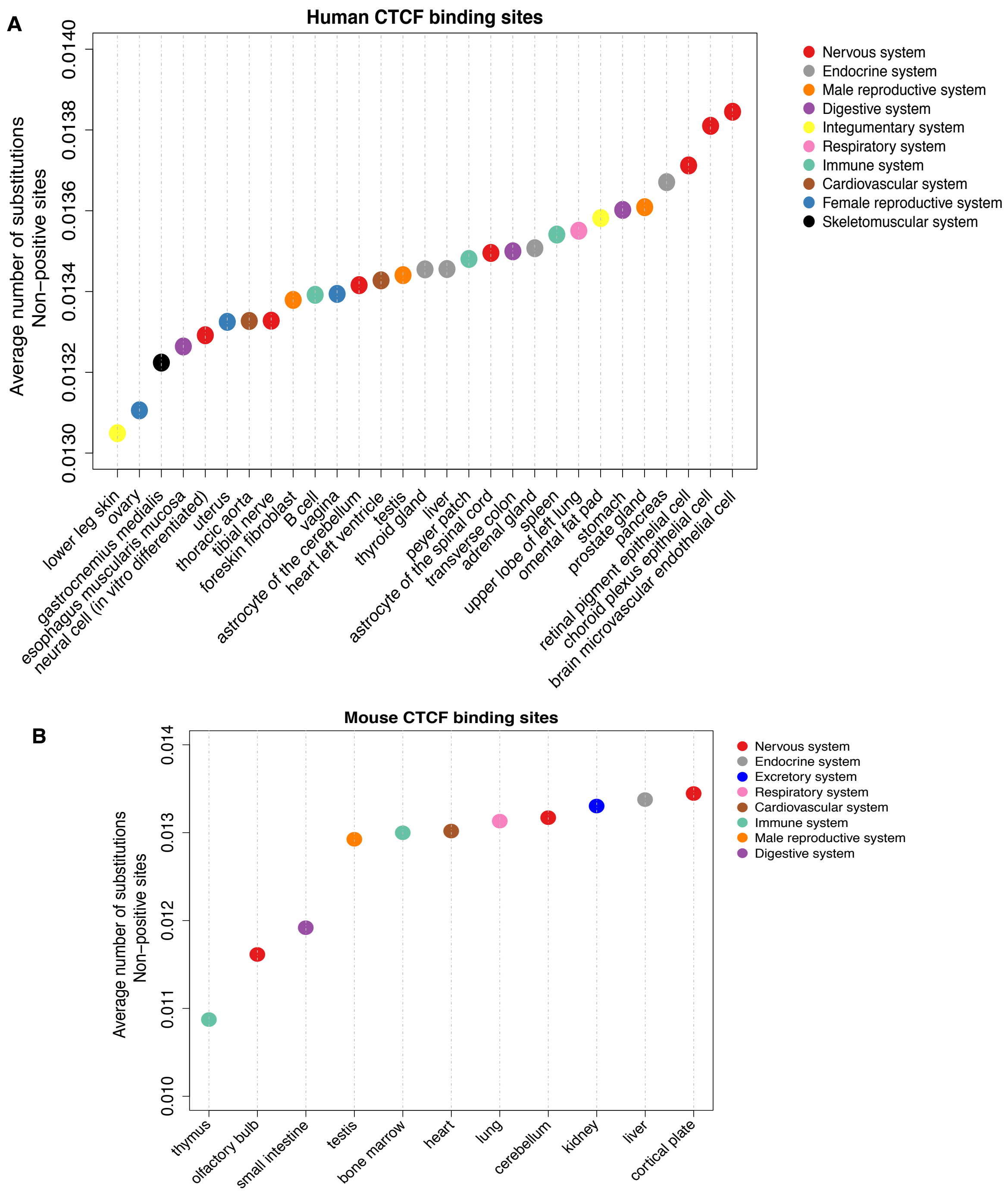


**Figure S19: Average number of substitutions for non-positive CTCF binding sites in different tissues/cell types.**

Non-positive binding sites are binding sites without evidence of positive selection (deltaSVM $p$-value >= 0.01). For human, the number of substitutions measured by the pairwise whole genome alignments between human and chimpanzee. For mouse, the number of substitutions measured by the pairwise whole genome alignments between C57BL/6J and CAST/EiJ. For each binding site, the number of substitutions was normalized by the length of the binding site.

A. CTCF binding sites in 29 human tissues/cell types.

B. CTCF binding sites in 11 mouse tissues.

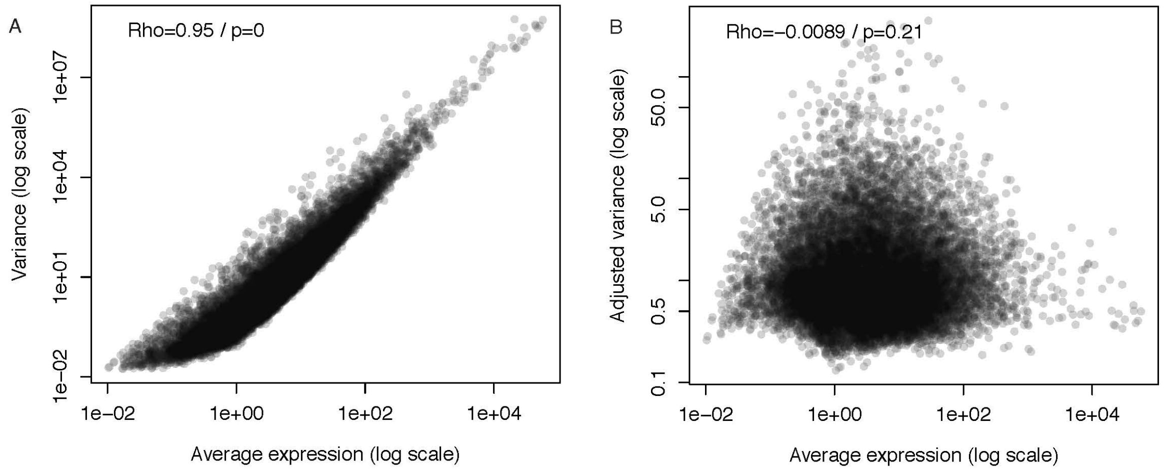


**Figure S20: Relationship between mean expression and variance.**

A. Spearman's correlation between mean expression and variance.

B. Spearman's correlation between mean expression and adjusted variance.

### 2 Supplementary tables

**Table S1: The number of peaks and average peak length for CEBPA, FOXA1 and HNF4A in three mouse species C57BL/6J, CAST/EiJ and SPRET/EiJ.**

| Factors / Species | CEBPA | FOXA1 | HNF4A |
|---|---|---|---|
| **C57BL/6J** | 41945 (270 bp) | 50683 (276 bp) | 68310 (298 bp) |
| **CAST/EiJ** | 51057 (266 bp) | 59780 (280 bp) | 72257 (289 bp) |
| **SPRET/EiJ** | 54035 (244 bp) | 64027 (271 bp) | 60142 (275 bp) |

**Table S2: The CTCF ChIP-seq datasets information for human and mouse.**

| Human | | | |
|---|---|---|---|
| tissue/cell type | Encode accession ID | # peaks | sample information |
| transverse colon | ENCFF547PLX | 42837 | male adult 37 years |
| stomach | ENCFF560VSK | 24358 | male adult 54 years |
| peyer patch | ENCFF777TZZ | 44111 | male adult 37 years |
| adrenal gland | ENCFF622LJD | 38269 | male adult 37 years |
| pancreas | ENCFF628TDS | 44068 | male adult 37 years |
| spleen | ENCFF459AHK | 20634 | male adult 37 years |
| upper lobe of left lung | ENCFF984EZB | 44154 | male adult 37 years |
| lower leg skin | ENCFF733VAF | 6905 | male adult 37 years |
| thyroid gland | ENCFF384ALJ | 40010 | male adult 54 years |
| gastrocnemius medialis | ENCFF101PDP | 23000 | male adult 54 years |

| | | | |
|---|---|---|---|
| heart left ventricle | ENCFF240UFV | 44400 | female adult 53 years |
| omental fat pad | ENCFF454PSG | 7257 | male adult 37 years |
| tibial nerve | ENCFF407FUL | 38250 | male adult 37 years |
| uterus | ENCFF282BOE | 34171 | female adult 53 years |
| ovary | ENCFF261BWI | 25291 | female adult 53 years |
| prostate gland | ENCFF899MQP | 42180 | male adult 37 years |
| testis | ENCFF432XLE | 24328 | male adult 37 years |
| liver | ENCFF690BYG | 38845 | male adult 32 years |
| B cell | ENCFF449NOT | 52144 | female adult 43 years |
| thoracic aorta | ENCFF330BPK | 36852 | male adult 37 years |
| astrocyte of the cerebellum | ENCFF660HHS | 45500 | primary cell |
| brain microvascular endothelial cell | ENCFF065LHJ | 61680 | primary cell |
| choroid plexus epithelial cell | ENCFF700ILD | 62000 | primary cell |
| astrocyte of the spinal cord | ENCFF312HCK | 47893 | primary cell |
| neuron (in vitro differentiated cells) | ENCFF618DDO | 51087 | in vitro differentiated cells |
| Esophagus muscularis mucosa | ENCFF735EHK | 31886 | male adult 32 years |
| retinal pigment epithelial cell | ENCFF139DOR | 55106 | primary cell |
| vagina | ENCFF176MPT | 26446 | female adult 53 years |
| foreskin fibroblast | ENCFF337WIE | 48000 | primary cell |
| Mouse | | | |
| tissue | Encode accession ID | # peaks | sample information |
| liver | ENCFF542WEE | 37167 | male adult (8 weeks) |
| lung | ENCFF605YVN | 29061 | male adult (8 weeks) |
| heart | ENCFF616HYA | 35386 | male adult (8 weeks) |
| kidney | ENCFF311HPG | 37370 | male adult (8 weeks) |
| bone marrow | ENCFF806PDR | 27977 | male adult (8 weeks) |
| cerebellum | ENCFF357KNB | 32463 | male adult (8 weeks) |
| cortical plate | ENCFF034VZI | 32919 | male adult (8 weeks) |

| | | | |
|---|---|---|---|
| olfactory bulb | ENCFF143RHK | 14427 | male adult (8 weeks) |
| small intestine | ENCFF319LOC | 28414 | male adult (8 weeks) |
| testis | ENCFF443TPY | 24188 | male adult (8 weeks) |
| thymus | ENCFF714WDP | 20199 | male adult (8 weeks) |
